# Benchmarking Geometric Deep Learning for Cortical Segmentation and Neurodevelopmental Phenotype Prediction

**DOI:** 10.1101/2021.12.01.470730

**Authors:** Abdulah Fawaz, Logan Z. J. Williams, Amir Alansary, Cher Bass, Karthik Gopinath, Mariana da Silva, Simon Dahan, Chris Adamson, Bonnie Alexander, Deanne Thompson, Gareth Ball, Christian Desrosiers, Hervé Lombaert, Daniel Rueckert, A. David Edwards, Emma C. Robinson

## Abstract

The emerging field of geometric deep learning extends the application of convolutional neural networks to irregular domains such as graphs, meshes and surfaces. Several recent studies have explored the potential for using these techniques to analyse and segment the cortical surface. However, there has been no comprehensive comparison of these approaches to one another, nor to existing Euclidean methods, to date. This paper benchmarks a collection of geometric and traditional deep learning models on phenotype prediction and segmentation of sphericalised neonatal cortical surface data, from the publicly available Developing Human Connectome Project (dHCP). Tasks include prediction of postmenstrual age at scan, gestational age at birth and segmentation of the cortical surface into anatomical regions defined by the M-CRIB-S atlas. Performance was assessed not only in terms of model precision, but also in terms of network dependence on image registration, and model interpretation via occlusion. Networks were trained both on sphericalised and anatomical cortical meshes. Findings suggest that the utility of geometric deep learning over traditional deep learning is highly task-specific, which has implications for the design of future deep learning models on the cortical surface. The code, and instructions for data access, are available from https://github.com/Abdulah-Fawaz/Benchmarking-Surface-DL.

## 1. Introduction

In recent years, deep learning methodologies have emerged as powerful tools for image analysis, displaying robustness to noise in structure, shape and pose. These methods have had profound success in the medical imaging field, becoming state-of-the-art for analysing datasets confounded by spatial variation and noise. Progress has been made in applications such as disease detection and segmentation of 2-dimensional (2D) and 3-dimensional (3D) images (Chen et al.,2018;Falk et al.,2019;McKinney et al.,2020;Havaei et al.,2017). However, it has been much more challenging to translate this same success to surface or mesh domains, such as those commonly used for cortical imaging (Dale et al.,1999;Fischl et al.,1999), since the mathematical foundations of established deep learning methods rely heavily on the highly regular and consistent structure of Euclidean grids (Cohen and Welling, 2016; Bronstein et al.,2017).

Geometric deep learning (gDL) is the branch of research tasked at adapting convolutional neural networks (CNNs) to irregular domains such as surfaces, meshes and graphs (Bronstein et al.,2017). As it remains an active area of research, it has not produced a single coherent framework for treating convolutions, but has instead led to numerous divergent directions, ranging from strict adherence to mathematical definitions of convolution, to some more flexible approaches. While this lack of consensus may be partly due to the immaturity of the field, the main consideration is that non-Euclidean data are not homogeneous, and encompass diverse data sources. These include graphs e.g. patient relational data (Fang et al.,2021;Choi et al.,2020), chemical structures (Fout et al.,2017) and surfaces, both regular (Perraudin et al.,2019) and irregular (Gainza et al.,2020), with each motivating its own research directions. Our primary motivation for this paper is to benchmark these methods within the context of neuroimaging analyses on the cortical surface. We are not aware of any prior existing review of this breadth in this particular context.

The human cortex displays significant heterogeneity of structural and functional organisation (Glasser et al.,2016;Frost and Goebel,2012;Fischl et al.,2008) and this challenges conventional approaches for analyses based on image registration (Coalson et al.,2018;Tucholka et al.,2012). Previous studies have repeatedly shown the advantage of modelling the cerebral cortex as a surface (Glasser et al.,2013;Fischl et al.,2008;Frost and Goebel,2012; Robinson et al.,2014, 2018;Van Essen et al.,2012;Yeo et al.,2009). However, analyses remain challenging since evidence points to the dissociation of cortical function and sulcal morphology (Fischl et al.,2008;Nenning et al., 2017;Robinson et al.,2018) as well as significant topographic variation of cortical organisation, even across healthy populations (Glasser et al.,2016; Gordon et al.,2017;Kong et al.,2019). Together these factors significantly limit the precision with which smooth and topologically-constrained techniques for spatial normalisation can support population-based comparison. The residual misalignments that result from this imprecise normalisation may then have an undesirable impact on downstream analyses, for example leading to ambiguous interpretation of functional connectivity (Bijsterbosch et al.,2018, 2019).

For this reason, we seek to evaluate the potential for gDL methods on sphericalised cortical surfaces to act as a registration-independent technique for analysing cortical imaging features, and set out to benchmark these against traditional techniques such as region-of-interest (ROI) analyses, as well as 3D CNNs trained on volumetric magnetic resonance images (MRI). We evaluate performance based on two types of tasks: segmentation and regression of cortical phenotypes, using data from the Developing Human Connectome Project (dHCP) (Hughes et al.,2017;Makropoulos et al.,2018; Bozek et al.,2018). This dataset is freely available to the academic community, and provides a range of cognitively and clinically relevant targets for prediction. Here, we choose prediction of postmenstrual age (PMA) at scan, and gestational age (GA) at birth: the former may be seen as a baseline task, since the phenotype is so strongly and clearly expressed across the dataset through rapid changes in cortical morphology and organisation; and the latter may be seen as a more challenging, but highly clinically important target, since it correlates with degree of prematurity and thus might support mechanistic explanations for poorer neurodevelopmental outcomes commonly seen following preterm birth (McBryde et al.,2020;Brydges et al., 2018;Twilhaar et al.,2018;Kovachy et al.,2015). We also investigate cortical parcellation (hereafter referred to as segmentation) into folding-based regions using the Melbourne Children’s Regional Infant Brain Surface (M-CRIB-S) atlas (Alexander et al.,2017, 2019b;Adamson et al.,2020), as this has been one of the most popular problems for cortical gDL schemes thus far (Zhao et al.,2019;Gopinath et al.,2019;Cucurull et al.,2018).

In what follows, we present a brief review of the literature, including the main variants of gDL and baseline models benchmarked by this paper. In Section 3 preprocessing steps, network architectures and experiments are laid out. Section 4 presents results, including a review of how interpretable and robust the networks are to different types of transformation. Finally, in Section 5, we discuss the implications of these findings for the future design of surface deep learning networks for cortical imaging.

## 2. Background and Related works

The CNN is the centrepiece of modern deep learning for image recognition and analysis. It is based on a key mathematical property of the convolution operation - equivariance to translation - where the output of a convolution shifts as the input is shifted. This property results in models capable of recognising objects irrespective of their location in an image, making networks parameter-efficient, practical to train, and to some extent location or registration-independent.

The challenge in mapping CNNs to non-Euclidean domains is that translations no longer represent meaningful transformations of the data. Rather, transformations of filters across surfaces are more suitably parametrised as rotations. Unfortunately, defining rotationally-equivariant surface convolutions is far from straightforward, and so existing methods usually involve some measure of compromise: either in terms their equivariance, computational efficiency or learning power. For a full explanation of the theoretical underpinnings of these constraints, please review the appendix (Appendix A).

In brief, geometric convolutions may be broadly classified into spatial or spectral methods, which reference the domain that the convolution is computed in. Of these, spatial methods (Zhao et al.,2019;Jiang et al., 2019;Monti et al.,2017) simulate the familiar concept of passing a localised filter over the surface. In practice, while expressive, such methods often approximate mathematically correct convolutions; since, due to lack of a single, fixed coordinate system it is not possible to slide a filter over a curved surface whilst maintaining consistent filter orientation (Figure A.1b). This means that spatial (otherwise known as template-matching) approaches are generally expressive, but not necessarily rotationally-equivariant by design.

Spectral methods, on the other hand, utilise an alternate representation in which the (generalised) Fourier transform of a convolution of two functions may be represented by the product of their Fourier transforms. This opens the door to alternate representations which estimate convolution from the spectra of general meshes or graphs (Bruna et al.,2013;Defferrard et al., 2016;Kipf and Welling,2016). Evaluating full spectral convolutions on a graph can be computationally expensive due to the repeated calculation of graph spectra and use of large non-local spectral filters, both of which scale as the square of the graph size. To reduce this computational cost and to bring graph convolution scaling on par with spatial convolutions, graph CNNs most often approximate spectral filters through polynomial approximation (see Defferrard et al. (2016) or Appendix A.0.2 for further details). These return simple, symmetric, centric filter patterns, which are naturally rotationally equivariant, but far less expressive than spatial counterparts. Fully expressive, and rotationally-equivariant spectral methods have been proposed for the sphere (S2CNN Cohen et al. (2018)) but these remain highly computationally intensive.

Understanding the downstream implications of these compromises in the context of cortical modelling is the prime motivating factor of this work. In the majority of cases, we will consider the cortical surface as a sphere and will benchmark gDL approaches either explicitly designed for spherical manifolds, or adaptable to them (Zhao et al.,2019;Jiang et al.,2019;Cohen et al., 2018;Kipf and Welling,2016;Defferrard et al.,2016;Monti et al.,2017). Representing the cortical surface as a sphere reflects standard practice within the field, specifically motivated by the fact that cortical shape is known to be a poor marker of cortical organisation, and hence many aspects of cognition and behaviour (Glasser et al.,2016;Fischl et al.,2008).

In this study, we hypothesise that explicit modelling of cortical shape is not necessary for prediction of developmental phenotypes. We test this hypothesis by benchmarking against two networks trained on representations of anatomical (white matter) surfaces: PointNet++ (Qi et al.,2017b), which models the cortical surface as point-clouds, in order to learn transform- and perturbation-invariant representations of shapes; and the Spectral-Matching Graph Convolutional Network (Spectral-Matching GCN) (Gopinath et al., 2019), which implements graph convolutional operations directly on native space data, aligned through graph spectral embeddings (Lombaert et al., 2013).

We separately investigate alternatives to gDL, such as simply converting the problem into a Euclidean one by unwrapping the spherical surface onto a 2D plane, use of 3D volumes, and benchmark against the common practice of hand engineering features by averaging cortical data within ROIs on the surface. A brief overview of the different models, their assumptions and their limitations is presented in Table 1.

**Table 1:** Methods Benchmarked (for more details see Appendix A)

| Method | Reference | Domain | Overview | Equivariant |
| --- | --- | --- | --- | --- |
| **Baseline approaches** |  |  |  |  |
| Projected ResNet/U-Net | He et al. (2016a)<br>Ronneberger et al. (2015) | 2D/3D grid | Euclidean CNN implemented for spherical data projected onto a 2D plane | No<sup>1</sup> |
| Volumetric ResNet/U-net | He et al. (2016a)<br>Ronneberger et al. (2015) | 2D/3D grid | 3D CNN implemented for volumetric T2w MRI data | Yes |
| PointNet++ | Qi et al. (2017a)<br>Qi et al. (2017b) | Point cloud | Transformation and perturbation-invariant, multi-resolution representations of point clouds | Yes |
| Spectral-Matching GCN | Gopinath et al. (2019) | Spectral embedding | Performs spectral matching (Lombaert et al., 2015) and then fits a mixture of Gaussians spatial filter in the spectral domain | Yes |
| **Spectral Methods** |  |  |  |  |
| S2CNN | (Cohen et al., 2018) | Sphere | Full spectral convolution on the sphere; computationally very expensive | Yes |
| ChebNet | (Defferrard et al., 2016) | General graph | Chebyshev polynomial based approximation of spectral filters | Yes |
| GConvNet | (Kipf and Welling, 2017) | General graph | Further simplifies ChebNet by using only filters of first-order (direct) neighbours | Yes |
| **Spatial Methods** |  |  |  |  |
| Spherical UNet | (Zhao et al., 2019, 2021) | Sphere | Fits a regularly tessellated, and consistently oriented hexagonal filter on top of an icosahedral spherical mesh | No |
| MoNet | (Monti et al., 2017) | General manifold | Fits adaptive spatial filters based on learning a mixture of isotropic and anisotropic Gaussians | Yes |
| UG-SCNN | (Jiang et al., 2019) | Sphere | Fits hand-engineered filters based on a finite difference approximation of gradient and Laplacian operators | No |

<sup>1</sup> Projection to the 2D plane results in non-uniform distortions

Recent applications of spatial techniques to cortical surface analysis include a number of frameworks trained on the task of cortical segmentation (Zhao et al.,2019;Zhao et al.,2021;Parvathaneni et al.,2019); networks trained to predict Alzheimer’s disease (Mostapha et al.,2018); approaches trained to perform sex classification (Seong et al.,2018), and a spherical gradient-based approach used to predict individualised task-contrasts from resting state functional MRI data (Ngo et al.,2020). While there are a broad range of approaches for approximating different shaped filters appropriate for the sphere (Zhao et al.,2019, 2021;Jiang et al.,2019;Seong et al., 2018), fitting spatial filters to more general surfaces can be challenging, and are mostly suited to smooth surfaces (Monti et al.,2017), not the highly convoluted surface of the cerebral cortex.

Spectral methods, on the other hand, define (approximated) convolutions through the generalised Graph Laplacian, allowing them to describe both structured and unstructured data, and adapt to many different surface shapes. Accordingly, graph spectral methods have again found applications in cortical segmentation (Gopinath et al.,2019;He et al.,2020;Cucurull et al., 2018), classification of Alzheimer’s disease and Autism Spectrum Disorder (Azcona et al., 2020), and for mapping cognitive function (Wu et al., 2020; Ribeiro et al., 2020; Liu et al., 2020). In other work, the rotation-equivariant and expressive spectral filters of S2CNN (Cohen et al., 2018) were used successfully to classify Alzheimer’s disease from mild cognitive impairment, using downsampled spherical cortical-imaging data (Barbaroux et al., 2020).

Note, while graph networks have also been used to make predictions from functional and structural connectivity data (Dsouza et al., 2021; Gadgil et al., 2020; Dahan et al., 2021; Kong et al., 2021; Ktena et al., 2018; Li and Duncan, 2020), and combine imaging with non-imaging data (Arya et al., 2020; Parisot et al., 2018; Wee et al., 2019); these latter problems are considered outside of the scope of this review.

This paper builds upon initial work by Vosylius et al. (2020), who compared point-cloud and mesh CNNs against GCNs and 3D CNNs on the task of predicting PMA at scan in the dHCP cohort. In this work, we compare a wider range of spatial and spectral gDL approaches, focus on implementing approaches for spherical manifolds, and benchmark on more tasks (predicting PMA at scan, GA at birth and cortical segmentation).

## 3. Materials and Methods

### 3.1. Acquisition Protocol

All experiments are run using neuroimaging data from the publicly available dHCP dataset^1^ (Hughes et al., 2017; Cordero-Grande et al., 2018; Kuklisova-Murgasova et al., 2012; Makropoulos et al., 2018; Schuh et al., 2017). This dataset consists of inversion recovery T1-weighted (T1w) MRIs (TR=4795 ms; TI=1740 ms; TE=8.7 ms; SENSE factor: axial=2.26, sagittal=2.66) with overlapping slices (resolution (mm) 0.8 × 0.8 × 1.6); and T2-weighted (T2w) MRIs (TR=12 s; TE=156 ms; SENSE factor: axial=2.11, sagittal=2.58) acquired with the same resolution. Both T1w and T2w scans were reconstructed (Cordero-Grande et al., 2018) and super-resolved (Kuklisova-Murgasova et al., 2012) to 0.5mm isotropic resolution acquired from preterm and term neonates spanning 24-45 weeks’ PMA. All neonates were scanned during natural sleep and informed written parental consent was obtained prior to imaging. The dHCP was approved by the National Research Ethics Committee (REC: 14/LO/1169).

### 3.2. Cohorts

Benchmarking was performed on three tasks: a) prediction of PMA at scan, b) prediction of GA at birth, and c) cortical segmentation. Table 2 and Figure 1 summarise the dataset profiles for the individual tasks. Some of the dHCP preterm neonates were scanned twice, once around birth and once at term-equivalent age. In order to predict scan age as a marker of healthy neurodevelopment, we exclude later preterm scans from the scan age experiment. For similar reasons, preterm first scans were excluded from the birth age experiment (to explore the impact of prematurity on term-equivalent age cortical development). Finally, the dataset used for cortical segmentation corresponds to a subset of term-age datasets for which M-CRIB-S (Adamson et al., 2020) cortical labels were available at the time of writing.

**Table 2:** Dataset Summary

| Dataset | Total | Prematurity |  | Sex |  |
| --- | --- | --- | --- | --- | --- |
|  |  | Preterm | Term | Male | Female |
| Postmenstrual age at scan | 530 | 111 | 419 | 288 | 242 |
| Gestational age at birth | 514 | 95 | 419 | 283 | 231 |
| Segmentation | 391 | 67 | 324 | 218 | 173 |

**Figure 1:**
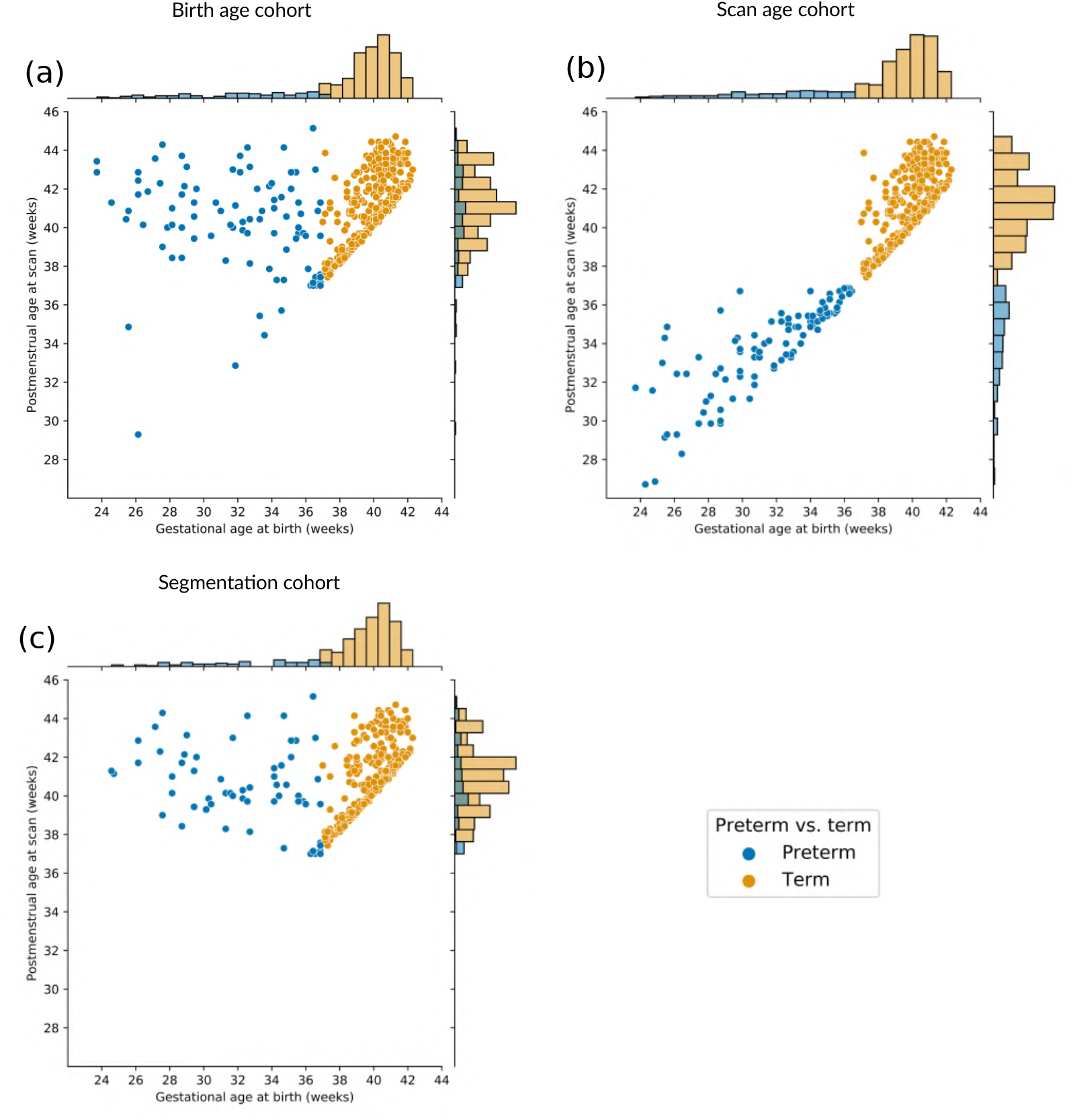
Gestational Age (GA) at birth and postmenstrual age (PMA) at scan of cohorts used for each benchmarking task. (a) predicting gestational age at birth, (b) predicting postmenstrual age at scan, (c) cortical segmentation.

### 3.3. Preprocessing

The full details of image reconstruction and preprocessing pipelines are described in Makropoulos et al. (2018) and references therein. In brief, motion corrected and reconstructed T2w and T1w images were passed through the dHCP structural pipeline^2^ (Makropoulos et al., 2018), which performed Draw-EM tissue segmentation (Makropoulos et al., 2014), surface extraction (Schuh et al., 2017) and inflation to return vertex-matched inner (white matter), outer (pial), midthickness, inflated and spherical surfaces. This process generated a number of univariate surface feature maps including: sulcal depth, mean curvature, cortical thickness (defined as the Euclidean distance between corresponding white matter and pial vertices) and T1w/T2w ratio maps, which are thought to reflect intracortical myelination (Glasser and Van Essen, 2011; Soun et al., 2017).

Segmentation labels corresponding to the M-CRIB-S atlas (Adamson et al., 2020) were also obtained. These represent a neonatal version of the Desikan-Killiany (Desikan et al., 2006) atlas; and were propagated to each subject using a combination of image registration and multi-atlas label fusion, implemented using a freely-available pipeline^3^ that leverages the popular FreeSurfer toolbox (Fischl, 2012).

All third-release data were registered to a modified version of the 40-week template of the dHCP spatiotemporal cortical surface atlas (Bozek et al., 2018), made to be left-right symmetric (Williams et al., 2021b). Whereas second release data, used for the cortical segmentation dataset, were registered to the original (non-symmetric) cortical surface template (Bozek et al., 2018). All registrations were performed using Multimodal Surface Matching (MSM) (Robinson et al., 2014, 2018) driven by sulcal depth as the sole feature. Both registered (template space) and unregistered (native space) data were used in this study (see Section 3.6.2). Prior to training (and testing) deep networks, surface feature maps were group-normalised to a mean and standard deviation of 0 and 1 respectively for each modality.

#### 3.3.1. Modelling the Cortex as an Icosphere

For all spherical network experiments, surface metric files were resampled from their native/template spheres to an icosphere of 40962 equally spaced vertices (sixth order icosphere; ico-6), using barycentric interpolation, implemented using Human Connectome Project (HCP) workbench software (Marcus et al., 2011). This icosahedral icosphere (Figure 2) represents a largely regular hexagonal tessellation of the sphere (with the exception of 12 pentagons). This supports straightforward down (and up) sampling, by virtue of the iterative process in which icospheres of different resolutions are generated (Figure 2); and, in the case of Spherical U-net (Zhao et al., 2019), the fitting of consistently shaped (hexagonal) spatial filters.

**Figure 2:**
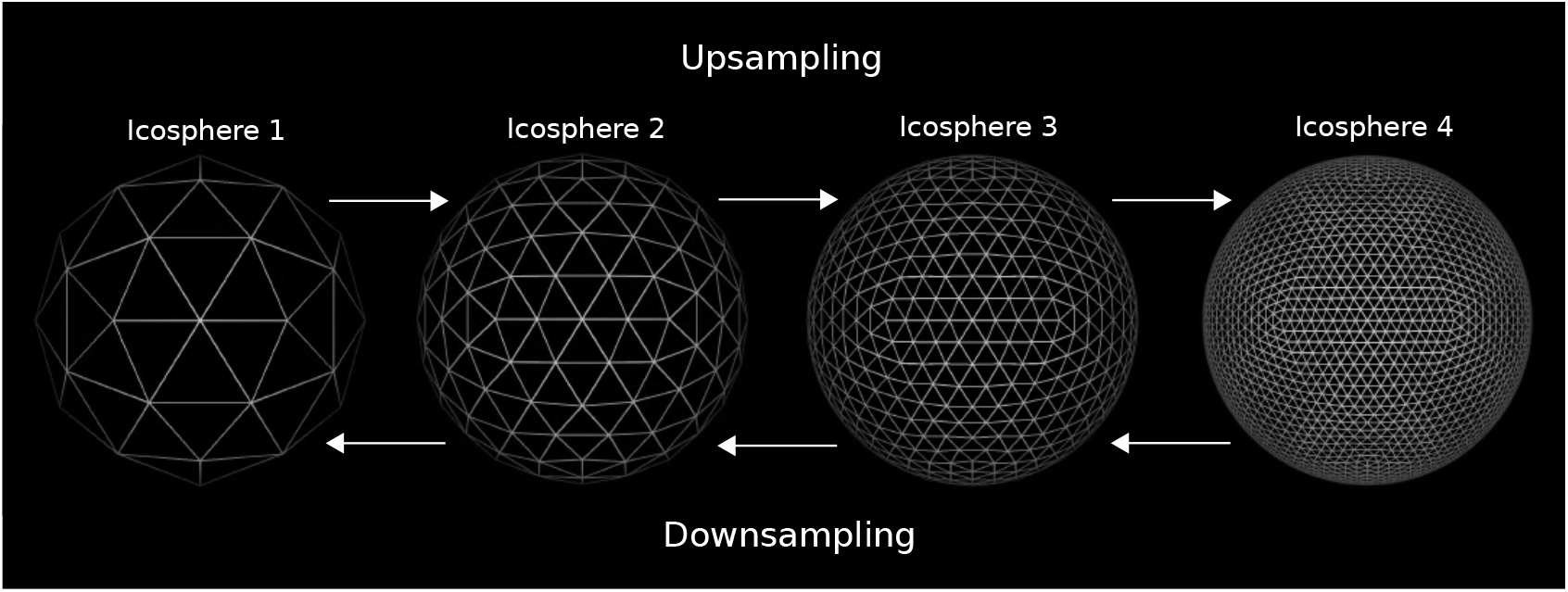
Icosahedrons can be efficiently up and downsampled to different resolutions.

#### 3.3.2. Method-Specific Image Preprocessing

Some of the networks benchmarked required additional or alternative preprocessing:

- **For the volumetric experiments**, T2w MRI scans were used. Native T2w images were registered to the 40-week template of the extended dHCP atlas (Schuh et al., 2018) using diffeomorphic multimodal (T1w/T2w) registration (Avants et al., 2011). To preserve age-related tissue maturation intensity differences, images were rescaled to [0,1] by normalising across the intensity range of the entire group. The images were brain extracted using blurred masks, and the extra-axial cerebrospinal fluid, ventricles and skull were removed in order to attend the model to brain tissue differences. Finally, the images were resized to 128 × 160 × 128 voxels. This setup reflects that used by Bass et al. (2021), which found that 3D CNNs did not train well on rigidly aligned volumetric dHCP data. One possible reason for this may be the difficulty in modelling both the shape *and* the rapid tissue maturational changes that occur across this relatively small cohort.
- **For 2D ResNet and S2CNN**, the icospheric surface data were projected directly from the icosphere to a 2D image of size 170 × 170 pixels using equirectangular projection (Figure 3vii). This projection was performed online using a precomputed sphere-to-plane projection map using linear interpolation for cortical feature maps across both regression and segmentation tasks, whilst nearest-neighbour interpolation was used only for labels in the segmentation task.

**Figure 3:**
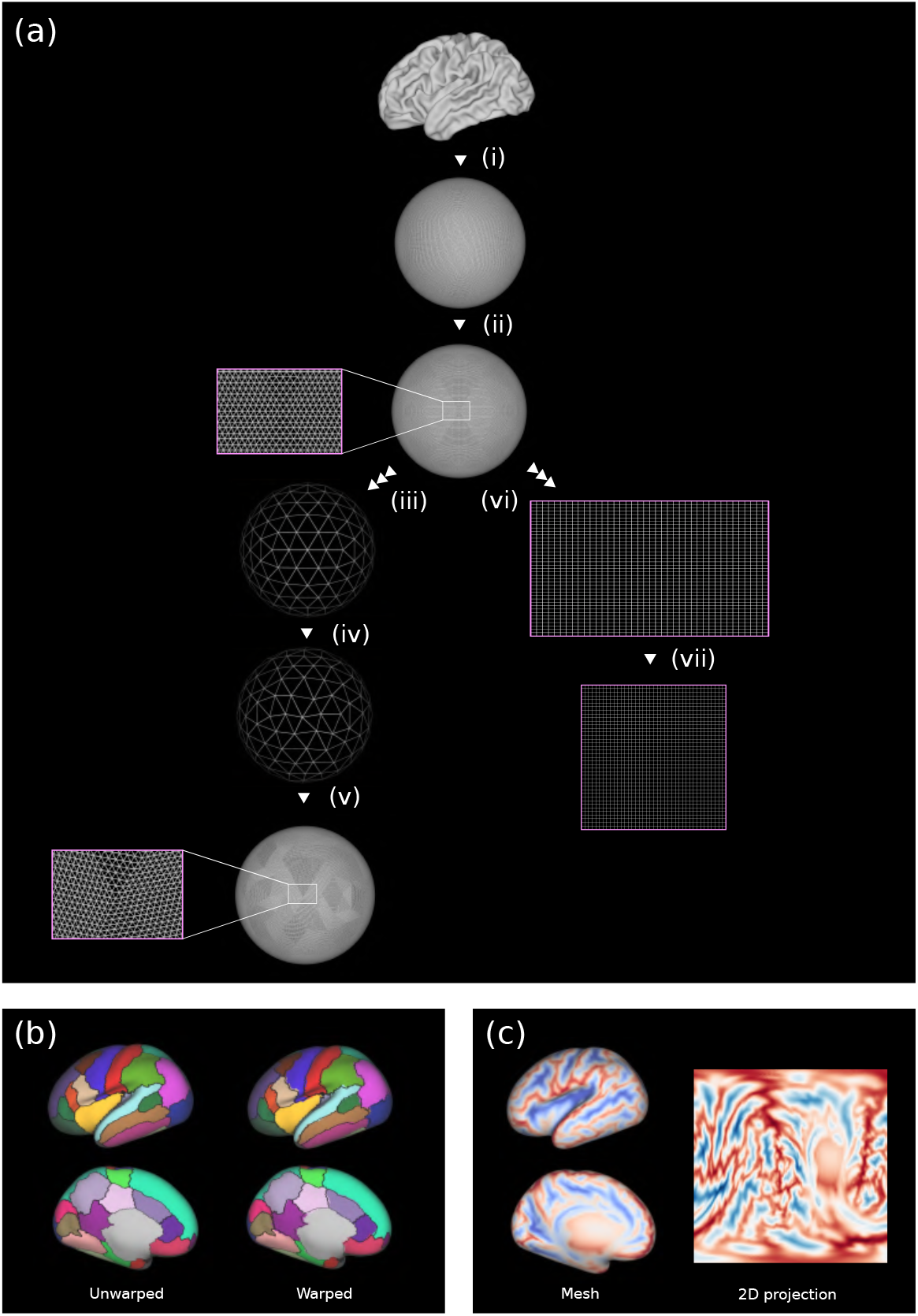
Data processing for benchmarking geometric deep learning methods. (a) Pipeline for generating data augmentations (only the left hemisphere is shown for illustrative purposes): (i) a 32k template white matter mesh is inflated and sphericalised, (ii) 32k spherical mesh is resampled to ico-6. To generate the 100 random warps, (iii) an Ico-6 icosphere was downsampled to a second order icosphere (Ico-2), (iv) vertices on the Ico-2 icosphere were randomly displaced to generate a random warp, (v) and this low-resolution warp was subsequently projected back to ico-6. Features corresponding to each example were then projected through this warp back down onto the regular ico-6 grid so as to generate a transformed image (not shown). To project data onto a 2-dimensional plane, (vi) data-points from the ico-6 icosphere were resampled to a precomputed grid, and (vii) this grid was reshaped to a size of 170 × 170. (b) shows an unwarped and warped single subject parcellations, respectively. The black outlines represent the borders of the non-warped parcellation. (c) shows a single subject sulcal depth map on a very inflated mesh and a 2-dimensional projection.
- **Cortical ROIs** were generated by first randomly segmenting the cortical surface into 150 Voronoi parcels, kept consistent across subjects. Surface feature maps (T1w/T2w ratio, curvature, cortical thickness and sulcal depth) were then averaged within each Voronoi parcel. Code for this is freely available^4^; for a visual example, please see Dimitrova et al. (2021b).
- **PointNet++** used anatomical white matter surfaces, resampled (following MSM registration) into vertex correspondence. In each case, left and right hemispheres were merged into one surface, and then the surface was then decimated to 20k nodes. In addition to vertex coordinates (Vosylius et al., 2020), PointNet++ also takes into account cortical metrics: T1w/T2w ratio, curvature, cortical thickness and sulcal depth.
- **Spectral-Embedding GCN** includes spectral embedding as part of its analysis pipeline; therefore, this method was run on native space white matter surfaces, with their original tessellation. Again the network was trained using the same cortical metrics (T1w/T2w ratio myelin maps, curvature, cortical thickness and sulcal depth) and spectral coordinates (Gopinath et al., 2019).

#### 3.3.3. Spherical Data Augmentation

Data were augmented through rotation and non-linear warping of the underlying icosahedral meshes. Rotations were defined as one of the 60 rotational symmetries of the icosahedron. In this way, rotations can be seen as a one-to-one mapping of vertices before and after rotation. These mappings were precomputed, and were used to perform rotations at train time. Of these 60 rotational configurations, 48 were used during training and 12 of them were set aside for testing. Data were also non-linearly warped to produce realistic variations in the data and improve network generalisation. Warped surface meshes (100 in total) were generated by randomly displacing the vertices on a second order icosphere (ico-2). The magnitude of this displacement was up to 1/8th of the distance between neighbouring points on the icosphere in order to retain point-point coherence. This warped ico-2 icosphere was resampled to an ico-6 icosphere using barycentric interpolation as described in Figure 3iv-v. For all models, other than those that also involve projection of data to a 2D plane, non-rigid warps were implemented offline to save run-time.

### 3.4. Architectures

Geometric deep learning comprises a very heterogeneous group of models (Bronstein et al., 2017), and benchmarking these comparably requires compromise between implementing models as they were designed, and fine-tuning model parameters to optimise performance. In this paper, the spherical network architectures were fixed for each class of task in order to more explicitly compare the impact of different choices of geometric convolution. The template architectures used for regression and segmentation are shown in Figure 4a and b respectively, with model specific parameters given in tables 3 and 4.

**Table 3:**
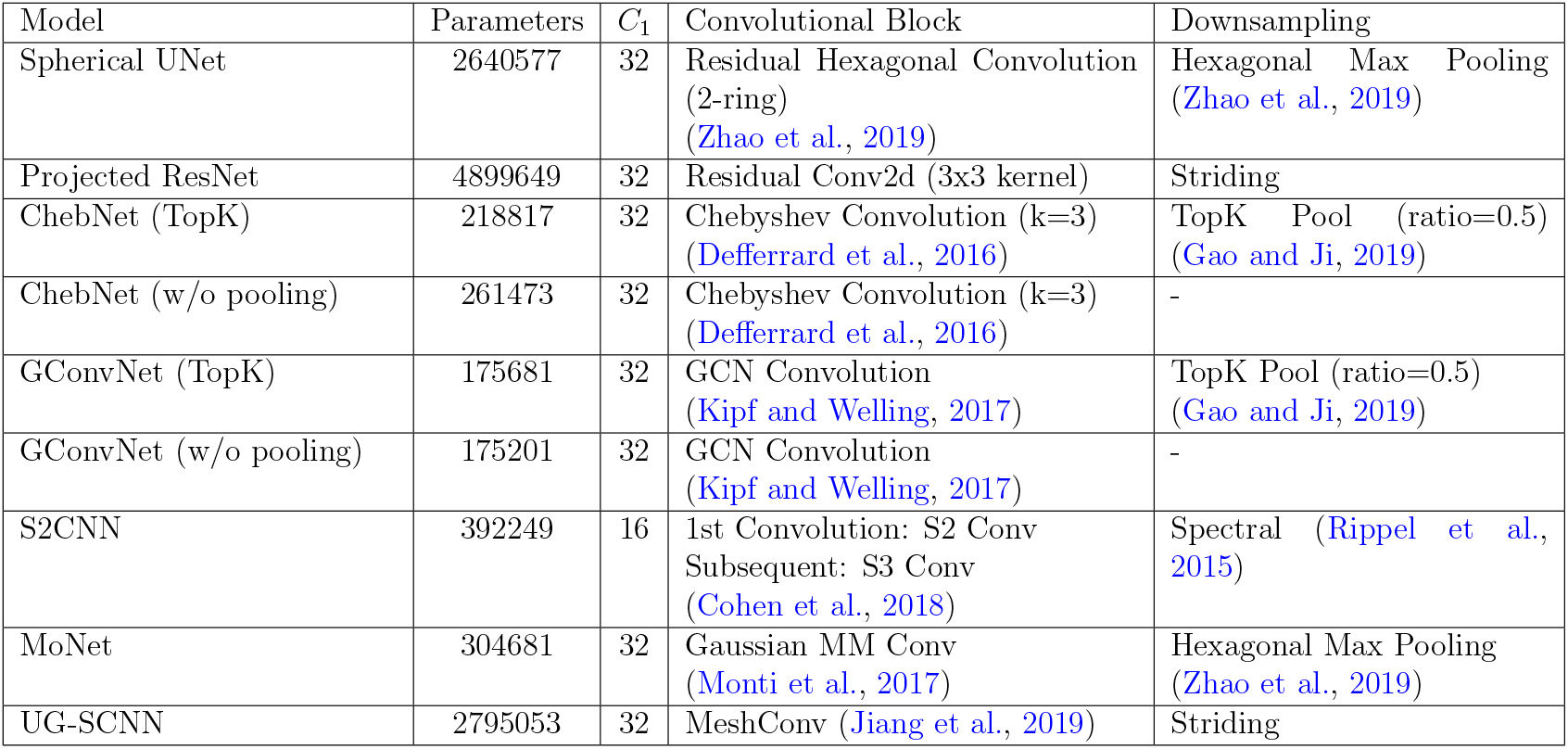
Architecture Summary: Regression

| Model | Parameters | $C_1$ | Convolutional Block | Downsampling |
| --- | --- | --- | --- | --- |
| Spherical UNet | 2640577 | 32 | Residual Hexagonal Convolution (2-ring)<br>(Zhao et al., 2019) | Hexagonal Max Pooling<br>(Zhao et al., 2019) |
| Projected ResNet | 4899649 | 32 | Residual Conv2d (3x3 kernel) | Striding |
| ChebNet (TopK) | 218817 | 32 | Chebyshev Convolution (k=3)<br>(Defferrard et al., 2016) | TopK Pool (ratio=0.5)<br>(Gao and Ji, 2019) |
| ChebNet (w/o pooling) | 261473 | 32 | Chebyshev Convolution (k=3)<br>(Defferrard et al., 2016) | - |
| GConvNet (TopK) | 175681 | 32 | GCN Convolution<br>(Kipf and Welling, 2017) | TopK Pool (ratio=0.5)<br>(Gao and Ji, 2019) |
| GConvNet (w/o pooling) | 175201 | 32 | GCN Convolution<br>(Kipf and Welling, 2017) | - |
| S2CNN | 392249 | 16 | 1st Convolution: S2 Conv<br>Subsequent: S3 Conv<br>(Cohen et al., 2018) | Spectral (Rippel et al., 2015) |
| MoNet | 304681 | 32 | Gaussian MM Conv<br>(Monti et al., 2017) | Hexagonal Max Pooling<br>(Zhao et al., 2019) |
| UG-SCNN | 2795053 | 32 | MeshConv (Jiang et al., 2019) | Striding |

**Table 4:** Architecture Summary: Segmentation

| Model | Parameters | $C_1$ | Convolutional Block | Downsampling | Upsampling |
| --- | --- | --- | --- | --- | --- |
| Spherical UNet | 16137701 | 32 | Residual Hexagonal Convolution (2-ring) (Zhao et al., 2019) | Hexagonal Max Pooling (Zhao et al., 2019) | Hexagonal Transposed Convolution |
| Projected ResNet | 8289349 | 32 | Residual Conv2d (3x3 kernel) | Mean 2d Pool (2x2) | Bilinear Interpolation |
| ChebNet (TopK) | 1054506 | 32 | Chebyshev Convolution (k=3) (Defferrard et al., 2016) | TopK Pool (ratio=0.5) (Gao and Ji, 2019) | TopK Unpooling |
| ChebNet | 2098433 | 32 | Chebyshev Convolution (k=3) (Defferrard et al., 2016) | Striding | Mean Unpool |
| GConvNet (TopK) | 353194 | 32 | GCN Convolution (Kipf and Welling, 2017) | TopK Pool (ratio=0.5) (Gao and Ji, 2019) | TopK Unpooling |
| GConvNet | 700825 | 32 | GCN Convolution (Kipf and Welling, 2017) | Striding | Mean Unpool |
| MoNet | 2797369 | 32 | Gaussian MMConv (Monti et al., 2017) | Hexagonal Max Pooling (Zhao et al., 2019) | Mean Unpool |
| UG-SCNN | 138547621 | 64 | MeshConv (Jiang et al., 2019) | Striding | Transposed MeshConv (Jiang et al., 2019) |

**Figure 4:**
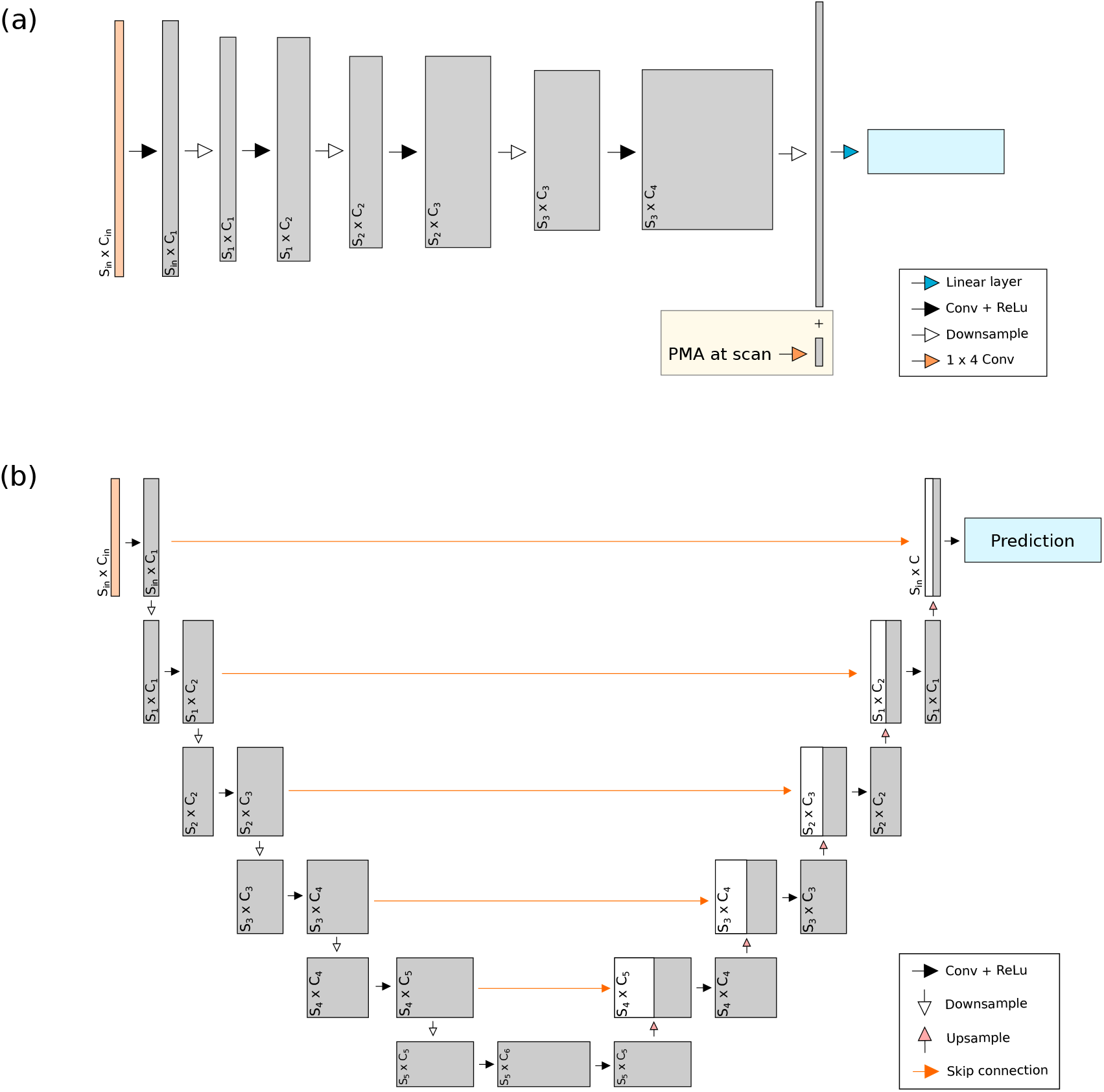
Model architectures: (a) regression, (b) segmentation

Generally, the regression models were run with a convolutional encoder with 4 layers of the appropriate convolution operation, each followed by a ReLU activation layer and a downsampling operation. Channel size doubled after each convolution from an initial value that was set to 32 for all models except S2CNN which began at 16 due to memory constraints. A fully connected layer was used to make a final age prediction. For birth age prediction, an additional 1D convolution was used to incorporate scan age as a confound. It was necessary to include this as the term-equivalent scans of each neonate were acquired for both preterm and term neonates, scanned at a range of ages (37 - 45 weeks’ PMA), meaning that scan appearance is dependent on both age at birth and at scan. The resulting output was concatenated with the encoded features prior to the linear layer, as shown in the figure.

Segmentation implemented a UNet (Ronneberger et al., 2015) like architecture, with a 6 layer encoding portion and a 6 layer decoder. Skip connections and upsampling operations were used as is standard practice for UNets, with a final softmax classification layer.

For Projected ResNet, striding was achieved on the 2D image domain in the usual way with a stride of 2. On the icosahedron, striding was used by UG-SCNN, ChebNet and GConvNet, and was achieved by forwarding only the vertices of the previous (smaller) icosahedral level. Hexagonal max pooling (Zhao et al., 2019), which takes the maximum of the direct neighbours of the vertices on the previous icosahedron level, was used by Spherical UNet and MoNet. S2CNN used spectral pooling - a simple reduction in the number of Fourier coefficients used to encode the data. For upsampling, Projected ResNet used a simple bilinear interpolation in 2D, Spherical UNet used a transposed convolution based on hexagonal neighbourhoods, and ChebNet, GConvNet, and MoNet all used a mean unpool operation on the icosahedron. This was achieved by simply averaging the two parent vertices of each new vertex under the icosahedron upsampling process (depicted in Figure 2). In addition, to the ChebNet and GconvNet architectures so far described, two (topK) variants were also run; where these were implemented utilising a learnable graph (un)pooling method known as TopK Pooling (Gao and Ji, 2019) (see Appendix A.0.3 for more details).

In terms of baseline comparisons: the 3D CNN trained on volumetric data used a ResNet structure starting with 1 standard convolutional layer (kernel=7, stride=2), followed by 3 ResNet layers (consisting of 2 ResNet blocks per layer (He et al., 2016b)), and 2 fully connected layers (with ReLU activation between them).

ROI-based prediction was estimated using random forests; 1000 trees were trained and 5-fold cross validation was used to optimise maximum tree depth and the maximum number of features considered per split. The optimal parameters for both tasks were tree depth equal to 3, and all features considered for each split.

The PointNet++ network architecture used is the same as in Vosylius et al. (2020), which consists of three hierarchical levels with a sampling layer, grouping layer, and PointNet layer. This is followed by a global max pooling to produce a vector of size 1024. Finally, three fully connected layers are used to generate either segmentations or predictions for age.

The Spectral Embedding GCN used 4 Gaussian Convolutional layers (the same convolutions used in MoNet) (Monti et al., 2017) with 6 kernels for both tasks, with an additional adaptive average pooling layer and three linear layers for regression. The spectral embedding preprocess generates an embedding using the normalized Graph Laplacian and aligns the resultant embeddings as detailed in Gopinath et al. (2019).

All networks were implemented in PyTorch with MoNet, ChebNet, GConvNet and PointNet++^5^ written using the PyTorch Geometric library (Fey and Lenssen,2019). The random forests code ^6^ (ROI model), S2CNN^7^, UG-SCNN^8^, Spherical UNet^9^ and Spectral Embedding GCN ^10^ were downloaded from existing open source repositories and, where necessary, networks were adapted for cortical surfaces (as described above). The code for the spectral alignment used in the spectral embedding GCN can be found at https://github.com/kharitz/aligned_spectral_embedding. All of the models and data used are available and can be implemented at https://github.com/Abdulah-Fawaz/Benchmarking-Surface-DL.

Note that most of the methods benchmarked were developed independently on non-neuroimaging tasks and are adapted from their original implementations to fit the domain, tasks used, and architectural templates found in this study. For many of the models this represents a significant departure from their original forms. However, the original names of the models were retained despite these changes and used across the different task-architectures. For example, Spherical UNet retains its name despite not being a UNet when applied to the age prediction tasks.

### 3.5. Implementation

All experiments were implemented on consistent train-validation-test split of 80-10-10, with batch sizes of 8 and 1 for regression and segmentation, respectively. All models were implemented on a Titan RTX 24GB GPU, and were trained and tested on a combined dataset of both left and right hemispheres, with right hemispheres mirrored to match left hemisphere orientation. During each epoch of training, 1 out of the 100 precomputed warped feature maps was randomly chosen per subject. If rotations were included in training, one of the 48 training rotations (or the identity transform) was then applied at random, online. No nonrigid transformations were applied during validation or testing.

In all cases optimisation of geometric networks was implemented with Adam learning rate = 0.001, betas = [0.9, 0.999], weight decay = 0.01 with mean square error (MSE) loss for regression, and standard (unweighted) Dice overlap ratio for segmentation. At each epoch, validation error was recorded and training was stopped after convergence - defined as 100 epochs without improvement on the best validation score. Regression experiments on spherical models were run four times with only the best performance over these runs quoted in the results for each model, alongside the standard deviation across all four runs. This was to compensate for the tendency of some of the models to occasionally converge at suboptimal local minima. The result reported for each run corresponds to testing on the check-pointed model with the best validation score. By contrast segmentation experiments were run only once.

The 3D CNN was trained using a smooth L1 loss, and the Adam optimiser with learning rate = 0.001, betas = [0.9, 0.999], and a weight decay = 0.001. The network was trained for 1000 epochs, with checkpointing, and the best validation model (i.e. best validation MAE) was used for testing. These are the same baseline model and hyper-parameters used in (Bass et al., 2021).

PointNet++ was trained using the same losses as the geometric networks, with batch sizes of 8 and 6 for regression and segmentation, respectively. Initial learning rates of 1*e*^−3^ and 5*e*^−3^ were used for prediction of PMA at scan and GA at birth experiments, respectively, along with a learning rate scheduler with a lower bound of 5*e*^−5^. Random rotations were applied during training, as is standard practice for point cloud-based models. An MSE loss criterion and Adam optimizer were used to train the model.

For the Spectral Embedding GCN, a fixed learning rate of 1e-3 was used with Adam optimizer, optimising for Cross Entropy + Dice loss for segmentation, and MSE for both regression of PMA at scan and GA at birth. No augmentation was applied during training.

### 3.6. Experiments

Models were benchmarked on three tasks: prediction of two developmental phenotypes (PMA at scan and GA at birth); and segmentation of the cortical surface. These benchmarks were performed under two different sets of experiments: the first investigating rotational equivariance of the models, and the second investigating model performance on native space data. Due to its large memory consumption, S2CNN was omitted from the segmentation experiments.

#### 3.6.1. Exploring Rotational Equivariance

The aim of these experiments was to explore the response of each model to unseen rotations. Of the 60 possible rotations, 48 of these were used during training with the remaining 12 only used during testing. In our experiments, the data were either unrotated (**U**), rotated by one of the 48 rotations used for training (**R**) or rotated by one of the 12 unseen rotations (**X**). We use the notation **TRAIN/TEST** to denote the experimental setup for training and testing e.g. **R/X** denotes a model was trained on the set of 48 rotations but tested on the set of 12 unseen rotations. By varying combinations of training and test rotations, the differences in model performance indicate the extent to which the models were rotationally equivariant. Specifically, the first experimental setup excluded rotations during training, but tested the trained models with (**U/R**) and without (**U/U**) the set of 48 rotations. These experiments addressed whether models generalised to rotations without any rotational augmentation. The second setup used the set of 48 rotations during training and tested without rotations (**R/U**), with the same 48 rotations (**R/R**), and on the 12 unseen test rotations (**R/X**). These experiments sought to determine the response of models to rotational augmentation during training i.e. whether the performance improved and whether models achieved rotational equivariance. For these experiments, we used data that were in template space.

#### 3.6.2. Training on native space data

In the second set of experiments, the above benchmarks were run on the same cortical metrics, but on each subjects’ own native space (unregistered) data. The aim was to investigate each models dependence on spatial normalisation. Performance was investigated with and without rotational augmentation during training, while, in all cases, non-rigid augmentations were used. Testing was performed solely on unrotated data.

### 3.7. Comparison of learning on Different Structures

We also compared the performance of spherical networks against networks implemented for direct analysis of native cortical anatomies, specifically PointNet++ (Qi et al., 2017b) and the Spectral-Embedding GCN method of Gopinath et al. (2019). In addition to shape, both methods were passed the same features used by the spherical methods: T1w/T2w ratio myelin maps, cortical thickness, curvature and sulcal depth.

### 3.8. Visualisation

For visualisation and interpretation, saliency maps were generated by the process of occlusion (Zeiler and Fergus, 2014; Huff et al., 2021). Specifically, hexagonal patches of the input cortical metrics were first replaced with uniform values; then these occluded images were passed through the trained models. Saliency maps were then generated as the change in the predicted output, with and without occlusion, for all patches, across the whole image. For surface data, occlusion was implemented for each individual modality separately and across all input channels simultaneously. In principle, occluding over individual modalities should give more interpretable saliency maps for those modalities, but can be unreliable if the same features are present across modalities at the same location. The occluded patches used were of size 276 vertices (corresponding to a 10-hop neighbourhood) with values set to 0. For the volumetric data, in which voxel intensity was the sole modality, occlusion was performed only once. The occluded patches were 10×10×10 voxels with values set to 0.5 (reflecting the different choice of normalization scheme used).

### 3.9. Data Access

Data from the dHCP project is openly released and is freely available^11^, subject to a data access agreement. In addition to files available from the public downloads, bespoke pre-processed data, ready for spherical network training and corresponding to the output of section 3.3, for both native and template space experiments, is accessible from a private GIN repo^12^ that will be made available to signatories of the dHCP terms of use. For more details on how to access this data, or to instead download scripts for pre-processing from scratch, please reference the code repository^13^.

## 4. Results and Discussion

### 4.1. Prediction of PMA at scan

The full results for the spherical (and baseline) experiments on prediction of PMA at scan can be found in Table 5 and we reference the columns therein. It can be seen from the **U/U** experiments that almost all of the models were capable of producing a mean absolute error (MAE) between 0.5 and 0.8 weeks for the prediction of PMA at scan, including the simple ROI baseline method. The performances were within the bounds of error for the measurement of GA from routine ultrasound, estimated to be ±5 − 7 days or 0.7 − 1 weeks (on Obstetric Practice et al., 2017). The exception was UG-SCNN, the worst performing model, with less than half the accuracy (1.24) of the baseline ROI method (0.63). UG-SCNN was by a significant margin the worst performing model across all experiments.

**Table 5:** Predicting PMA at scan, MAE in weeks. **A/B** indicates the type of rotation during training and testing respectively. **U** represents no rotations, **R** represents seen rotations and **X** represents unseen rotations. The first figures in the tables represent the best MAE, with the figures in brackets representing the standard deviation.

| Method | Network | U/U | U/R | R/U | R/R | R/X |
| --- | --- | --- | --- | --- | --- | --- |
| Baseline | Projected ResNet | 0.58 (0.16) | 9.4 (0.68) | 0.71 (0.10) | 0.70 (0.10) | 0.94 (0.04) |
|  | ROI | 0.63 (0.53) | - | - | - | - |
|  | Volumetric (3D ResNet) | 0.67 | - | - | - | - |
| Spectral | S2CNN | 0.63 (0.02) | **0.65** (0.02) | 0.63 (0.02) | 0.65 (0.02) | 0.69 (0.02) |
|  | ChebNet (TopK) | 0.67 (0.29) | 0.66 (0.33) | 0.71 (0.14) | 0.70 (0.12) | 0.70 (0.13) |
|  | ChebNet (w/o pooling) | 0.59 (0.37) | 0.66 (0.40) | 0.64 (0.01) | 0.64 (0.01) | 0.63 (0.01) |
|  | GConvNet (TopK) | 0.80 (0.27) | 0.78 (0.27) | 0.65 (0.004) | 0.66 (0.05) | 0.67 (0.04) |
|  | GConvNet (w/o pooling) | 0.75 (0.13) | 0.74 (0.12) | 0.73 (0.02) | 0.71 (0.01) | 0.71 (0.02) |
| Spatial | Spherical UNet | **0.57** (0.18) | 20.2 (2.5) | 0.87 (1.44) | 0.88 (2.54) | 2.49 (1.45) |
|  | MoNet | **0.57** (0.02) | 0.76 (0.07) | **0.57** (0.03) | **0.57** (0.02) | **0.57** (0.03) |
|  | UG-SCNN | 1.24 (0.87) | 8.16 (3.70) | 3.17 (2.10) | 4.57 (2.43) | 4.90 (2.48) |

Excluding UG-SCNN, the performance of all models for this problem under these conditions was high; however, the problem of predicting PMA at scan is not a difficult one. Overall, Spherical UNet and MoNet were the best performing models (both at 0.57 MAE) followed closely by Projected ResNet (0.58) and ChebNet (without pooling) (0.59). The second-worst performing model, GConvNet (TopK) achieved an MAE of 0.80 - corresponding to a difference in accuracy between the best and worst models of just over two days on average, which lies within measurement error. The baseline 3D volumetric ResNet achieved an MAE of 0.67 weeks, an average performance amongst the models.

Comparing the results of the **U/U** experiments to the **U/R** experiments, some models differed significantly in performance on the rotated test set, whereas others exhibited insignificant change. These differences reflect the absence or presence of rotational equivariance. In particular, Spherical UNet, UG-SCNN and Projected ResNet experienced drastic drops in performance, with accuracies within a week falling to over a month, reflecting that they are not rotationally equivariant (NRE) - exactly as predicted by theory. In contrast, the models expected to be rotationally equivariant (RE) (S2CNN, ChebNet, GconvNet and MoNet) had no significant change in performance, although MoNet had a moderate increase of 0.19 weeks (∼ 1.4 days) - significantly less than the large increases (8+ weeks) seen in the NRE models, but more than the other RE models which changed by less than half a day. Comparing the **R/U** and the **U/U** experiments showed another difference between the NRE and the RE models. The RE models showed no significant degradation in performance when rotations were added during training (change *<* 0.1) with some, such as GConvNet (TopK), even improving in performance by about a day on average. In contrast, all of the NRE models exhibited some drop in performance, ranging from 0.14 weeks for Projected ResNet to higher values of 0.3 and 1.9 weeks for Spherical UNet and UG-SCNN respectively. This too reflects theory as the convolution operations of the NRE models were not able to capture rotational variations naturally, and instead had to learn different rotational configurations separately. The differences in magnitude of this degradation may correspond to the varying degree in which the networks are able to learn multiple rotations. Comparing **R/U** to **R/R** and noting the very small change in performance for both the NRE and RE models, it might appear that adding rotations during training allowed the NRE models to generalise to rotated data. However, there is a distinction between seen and unseen rotations, and comparing across the **R/U, R/R** and **R/X** columns reveals the same divergence between the NRE and RE models, with the RE models showing little change in performance across all three testing groups, including on unseen rotations (**R/X**) with an overall MAE change *<* 0.1 across all setups; whereas the NRE models showed generalisation to seen rotations (**R/R**) but a drop in performance on unseen rotations (**R/X**). This is an important result as it shows that rotational equivariance cannot be obtained by the addition of rotations as an augmentation during training. Nevertheless, this drop in performance was less drastic than the previous experiments, and so some degree of robustness to rotations has been learned.

With respect to the rotational equivariance of the Projected ResNet baseline, this showed a greater degree of generalisation to rotations than the other NRE models, with a much smaller overall variation in performance (around 0.3 overall) when trained with rotations. This may be attributed to either the power of the model itself, or the simplicity of the scan age prediction task.

Overall the best model, under augmentations, with 0.57 MAE across all (R/U, R/R and R/X) configurations was MoNet. The fact that it shows no drop in performance when tested on unseen rotations, demonstrates that, under suitable training conditions, it is indeed rotationally equivariant, as indicated in Table 1.

### 4.2. Prediction of GA at birth

Results of predicting GA at birth are shown in Table 6. As a correlate of prematurity but confounded by PMA at scan, this makes for a more difficult task. As a result, all experiments returned a higher MAE, with the ROI method performing worst, with an MAE of 4.0 weeks, arguably justifying the use of deep learning on this task. From the **U/U** column of the table, it can be seen that UG-SCNN was again the worst performing gDL model with an MAE of 3.55 weeks. The 3D Residual Network on the volume again achieved a median performance among the models with an MAE of 1.84 weeks, indicating that whilst performing deep learning directly on the volume can produce good results, performing deep learning on the surface may be seen as equal or better.

**Table 6:** Predicting GA at birth, MAE in weeks. **A/B** indicates the type of rotation during training and testing respectively. **U** represents no rotations, **R** represents seen rotations and **X** represents unseen rotations. The first figures in the tables represent the best MAE, with the figures in brackets representing the standard deviation.

| Method | Network | U/U | U/R | R/U | R/R | R/X |
| --- | --- | --- | --- | --- | --- | --- |
| Baseline | Projected ResNet | 1.33 (0.02) | 12.83 (3.08) | 1.63 (0.01) | 1.37 (0.07) | 2.63 (0.31) |
|  | ROI | 4.00 (1.83) | - | - | - | - |
|  | Volumetric (3D ResNet) | 1.84 | - | - | - | - |
| Spectral | S2CNN | 1.35 (0.68) | **1.40** (0.67) | 1.55 (0.46) | 1.64 (0.44) | 1.61 (0.45) |
|  | ChebNet (TopK) | 2.29 (0.10) | 2.20 (0.14) | 2.28 (0.15) | 2.22 (0.11) | 2.24 (0.06) |
|  | ChebNet (w/o pooling) | 1.57 (0.15) | 1.73 (0.15) | 1.63 (0.15) | 1.73 (0.10) | 1.74 (0.08) |
|  | GConvNet (TopK) | 1.90 (0.47) | 1.94 (0.47) | 2.03 (0.12) | 2.02 (0.16) | 2.00 (0.18) |
|  | GConvNet (w/o pooling) | 1.77 (0.26) | 1.86 (0.23) | 1.52 (0.39) | 1.50 (0.37) | 1.54 (0.32) |
| Spatial | Spherical UNet | **0.85** (0.17) | 5.89 (7.6) | 2.19 (0.38) | 3.05 (1.41) | 3.50 (0.48) |
|  | MoNet | 1.44 (0.08) | 2.33 (0.72) | **1.33** (0.21) | **1.33** (0.22) | **1.31** (0.25) |
|  | UG-SCNN | 3.55 (1.83) | 3.55 (1.94) | 2.81 (2.49) | 2.80 (2.47) | 2.95 (2.62) |

Overall, Spherical UNet performed the best by some margin with an MAE of 0.85 weeks, followed by Projected ResNet and S2CNN, with an MAE of 1.33 weeks and 1.35 weeks, respectively. It is noteworthy that these are the three methods with fully expressive filters. These were followed closely by MoNet (1.44 weeks) and ChebNet without pooling (1.57 weeks), then GConvNet without pooling (1.77 weeks). The performance of both ChebNet and GConvNet was consistently better without pooling than with TopK pooling, a trend consistent across all of the experiments.

Comparing **U/U** to **U/R**, the same trends appeared as for scan age prediction. The RE methods did not significantly change in performance when tested on rotated data whereas the NRE methods (Projected ResNet and Spherical UNet) dropped significantly, once again indicating the differences in rotational equivariance between these models. From the **U/U** results to **R/U**, the performance of Spherical UNet dropped significantly when rotations were added to training, from the best performing model to one of the worst. The RE models experienced only small changes with most improving slightly. This difference indicates that the RE models were able to capture variations present in the dataset augmented by rotations.

The difference in consistency between the NRE and RE models when tested on seen and unseen rotations was more pronounced than for the scan age experiments. This is highlighted by the range of accuracies shown over the **R/U, R/R** and **R/X** experiments. The NRE models, including Projected ResNet, showed a wider range in performance with clear drops in accuracy when tested on unseen rotations, whilst the RE models did not change significantly across all test sets. This confirms the findings on scan age that NRE models cannot be made RE by augmenting training with rotations.

Although UG-SCNN gave the illusion of rotational equivariance, it only predicted a narrow range of birth ages (between 36 and 38 weeks) around the mean birth age (true range between 27 and 44 weeks), and therefore failed to accurately predict GA at birth.

### 4.3. Segmentation

The results for segmentation of the cortical surface can be found in Table 7, with an example of predicted cortical segmentations, for each model, shown in Figure 5. As M-CRIB-S segmentation may be seen as resulting from a combination of folding based image registration followed by multi-atlas label fusion (to address individual folding variability) the template-space segmentation experiments shown here may be seen as validating the potential of deep networks to learn this residual variation.

**Table 7:** Mean Dice overlap ratio for cortical segmentation data in template space and standard deviation across all subjects.

| Method Type | Network | U/U | U/R | R/U | R/R | R/X |
| --- | --- | --- | --- | --- | --- | --- |
| BaseLine | Projected UNet | $0.936 \pm 0.011$ | $0.187 \pm 0.132$ | $0.912 \pm 0.040$ | $0.889 \pm 0.068$ | $0.709 \pm 0.096$ |
| Spectral | ChebNet (TopK) | $0.319 \pm 0.037$ | $0.305 \pm 0.037$ | $0.309 \pm 0.049$ | $0.310 \pm 0.049$ | $0.310 \pm 0.050$ |
| | ChebNet (w/o TopK) | $0.939 \pm 0.008$ | $0.298 \pm 0.137$ | $0.929 \pm 0.014$ | $0.933 \pm 0.011$ | $0.923 \pm 0.016$ |
| | GConvNet (TopK) | $0.246 \pm 0.052$ | $0.245 \pm 0.052$ | $0.242 \pm 0.050$ | $0.242 \pm 0.050$ | $0.242 \pm 0.050$ |
| | GConvNet (w/o TopK) | $0.930 \pm 0.010$ | **$0.637 \pm 0.063$** | $0.929 \pm 0.009$ | $0.932 \pm 0.008$ | $0.922 \pm 0.010$ |
| Spatial | Spherical UNet | $0.941 \pm 0.008$ | $0.181 \pm 0.127$ | $0.940 \pm 0.007$ | $0.939 \pm 0.007$ | $0.191 \pm 0.056$ |
| | MoNet | **$0.946 \pm 0.007$** | $0.273 \pm 0.180$ | **$0.944 \pm 0.008$** | **$0.945 \pm 0.008$** | **$0.942 \pm 0.009$** |
| | UG-SCNN | $0.887 \pm 0.030$ | $0.150 \pm 0.126$ | $0.292 \pm 0.038$ | $0.246 \pm 0.036$ | $0.238 \pm 0.031$ |

**Figure 5:**
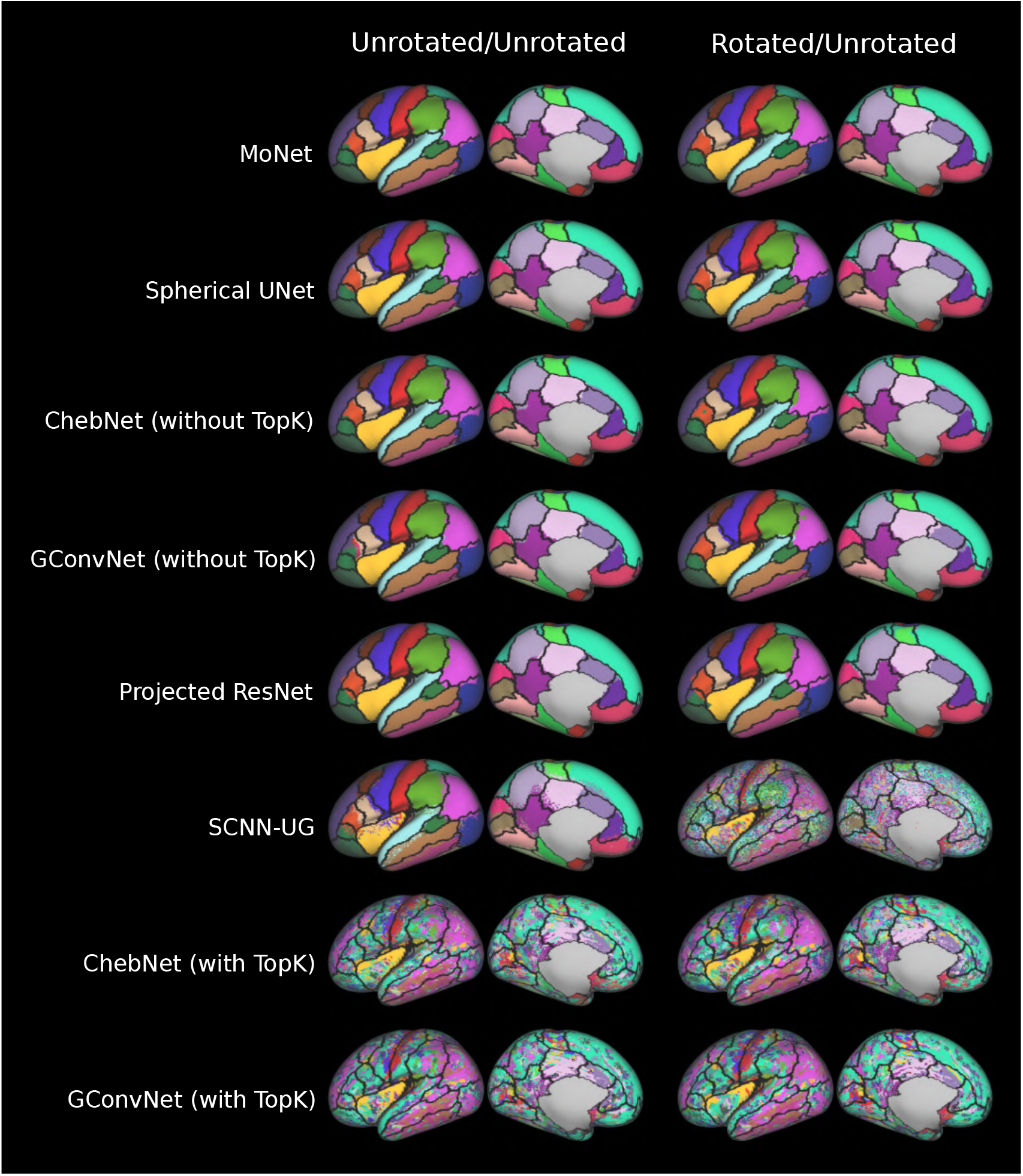
Example of predicted cortical surface segmentations for a single subject across all models for U/U and R/U experiments in template space. Black lines show ground truth labels from M-CRIB-S pipeline. Results are shown on a 40 week PMA very inflated left hemispheric surface.

From the **U/U** experiments, most of the models performed extremely well when trained and tested without rotational augmentations, with mean Dice overlap ratios between 0.930 and 0.946. However, models with TopK pooling completely failed at the segmentation task, likely due to the inability of the TopK pooling to accurately consolidate spatial information (Figure 5). In contrast to its performance on phenotype prediction, UG-SCNN performed relatively well with a mean Dice overlap ratio of 0.887, although the segmentations contained high levels of noise resulting in spatially non-contiguous parcels. As with prediction of PMA at scan, the top two performing models on segmentation were MoNet and Spherical UNet.

Comparing **U/U** to **U/R** revealed the same expected trend that the RE models generalised better to unseen rotations than the NRE models. Interestingly, unlike prediction of scan age and birth age, the RE models experienced a significant drop in performance on unseen rotated data, albeit less than the NRE models. The NRE models achieved Dice overlap ratios *<* 0.2 on test rotations, whereas the RE models dropped to between 0.2 and 0.3. The exception was GConvNet without TopK pooling which retained a Dice overlap ratio of 0.637. These results indicate that models memorised the data rather than learning generalisable filters, a conclusion that is supported when rotations are added to training.

From the results of the **R/U** and **U/U** experiments, UG-SCNN failed to accurately capture different rotational configurations and became one of the worst performing models. Projected ResNet dropped in performance by approximately 0.1 but Spherical UNet retained its performance in a manner similar to the RE models. Comparing **R/X** and **R/R** to **U/R**, however, reveals that adding rotations to training resulted in more robust generalisation to both seen and unseen rotations for the RE models, all of which gave uniform Dice overlap ratios across all test sets (change *<* 0.01). Conversely, the NRE models showed the same significant trends as found for the age prediction tasks, with some robustness to seen rotations (**R/R**) but significant degradation for unseen rotations (**R/X**), with Dice overlap ratios of 0.709 and 0.191 for Projected ResNet and Spherical UNet, respectively.

We also analysed the performance of each model, at a regional level (Figures 6 and 7) - this time only considering MoNet, Spherical UNet, ChebNet and GConvNet (without TopK pooling), and Projected ResNet. Here, the performance of each model, in terms the stability of the prediction across all test examples, was explored. Overall, larger regions such as the superior frontal gyrus and insula had higher mean and lower standard deviation Dice overlap (across test subjects), whereas smaller regions such as the bank of the superior temporal sulcus and frontal pole performed more variably. These findings are consistent with other studies focused on cortical segmentation in neonates (Adamson et al., 2020; Zhao et al., 2019) and adults (Gopinath et al., 2019; Parvathaneni et al., 2019; Williams et al., 2021a), and partly reflect an inherent limitation of using the Dice overlap ratio as a performance measure for multiclass segmentation (Reinke et al., 2021). However, arbitrary anatomical boundaries (especially in the case of the bank of the superior temporal sulcus) may also explain lower mean and higher standard deviation Dice overlap ratios (Adamson et al., 2020; Klein and Tourville, 2012).

**Figure 6:**
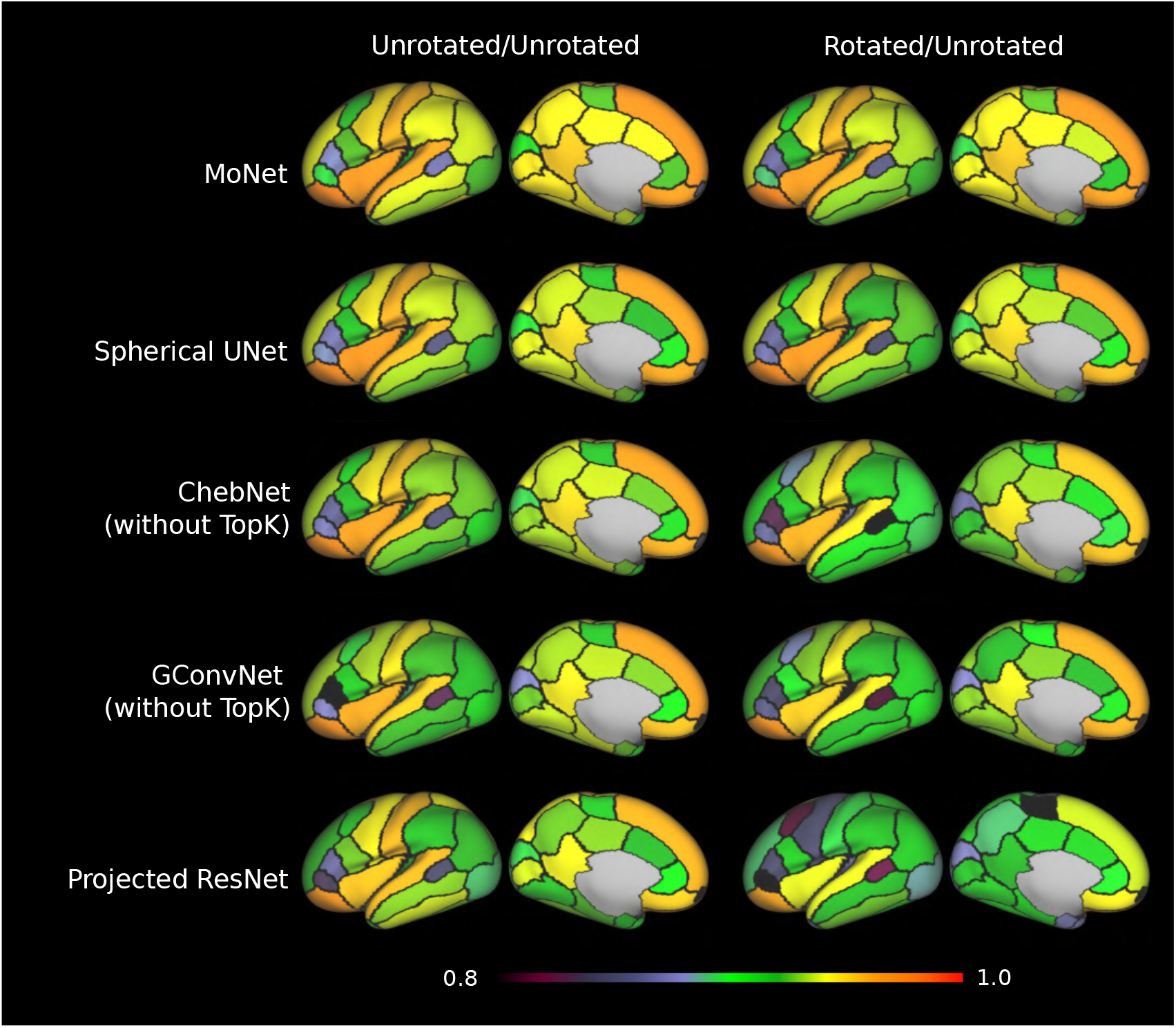
Mean Dice overlap ratio per region, for U/U (column 1) and R/U experiments (column 2). Dice overlap ratio calculated across both hemispheres. Black outlines represent group average cortical segmentation. Results shown on a 40 week PMA very inflated left hemispheric surface.

**Figure 7:**
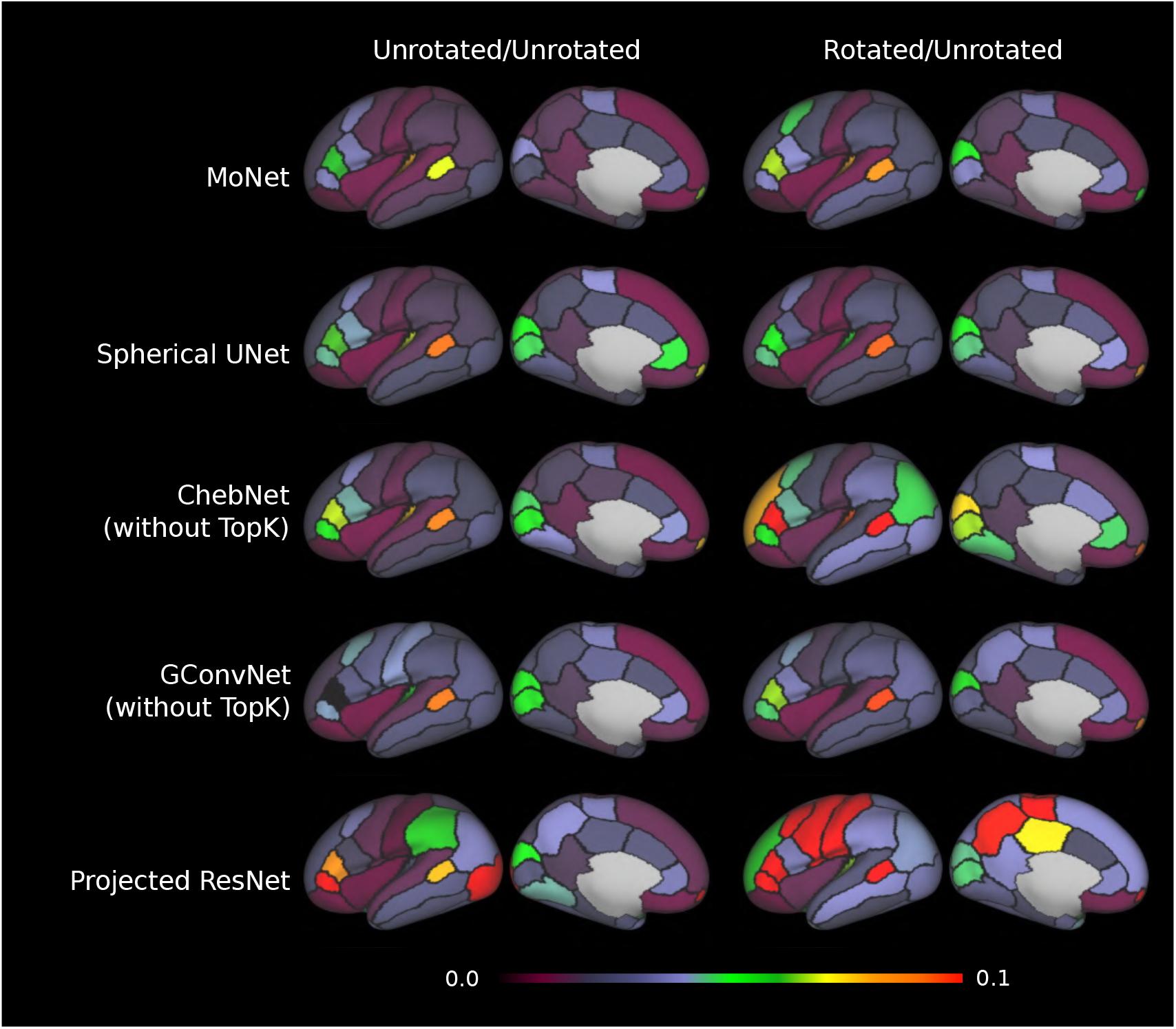
Standard deviation Dice overlap ratio, for U/U and R/U experiments. Dice overlap ratio calculated across both hemispheres. Black outlines represent group average cortical segmentation. Results shown on a 40 week PMA very inflated left hemispheric surface.

An exception to this trend was the regional performance of Projected ResNet (**R/U**), where the precentral gyrus and caudal middle frontal gyrus exhibited lower mean and higher standard deviation Dice overlap. One possible explanation for this is that these regions become greatly distorted as a result of projection to a 2D plane (see Figure 3). This highlights the importance of regional analysis during benchmarking of cortical segmentation models as summarising performance in a single global measure can mask important model limitations.

### 4.4. Performance on Native Space Data

Performances of each network on native space data are shown in Tables 8 to 10. Before describing the results, it is worth noting how data in native space differs from that in template space. Figure 8 shows the inter-subject variability of two regions (postcentral and inferior temporal gyri) in native space, visualised on the template inflated surface. It can be seen that there is significant consistency in both the shape and position of each region across subjects, due to the fact that they are already volumetrically rigidly aligned before surface extraction. However, subjects still differ by a global rotation and a non-linear deformation, and all subjects are rotated out of correspondence with the template (for instance, the inferior temporal gyrus in native space overlaps the superior and middle temporal gyri in template space).

**Table 8:** Predicting PMA at scan, MAE in weeks on native space data. Columns refer to the presence or lack of rotational augmentations during training. The first figures in the tables represent the best MAE, with the figures in brackets representing the standard deviation.

| Method | Network | Unrotated | Rotated |
| --- | --- | --- | --- |
| Baseline | Projected ResNet | 0.68 (0.34) | 1.13 (0.27) |
| Spectral | S2CNN | 0.73 (0.25) | 0.79 (0.2) |
|  | ChebNet (TopK) | 0.67 (0.42) | 0.66 (0.63) |
|  | ChebNet (w/o pooling) | 0.77 (0.49) | 0.68 (0.41) |
|  | GConvNet (TopK) | 0.70 (0.38) | 0.73 (0.28) |
|  | GConvNet (w/o pooling) | 0.75 (0.26) | 0.70 (0.46) |
| Spatial | Spherical UNet | 0.87 (0.50) | 2.90 (0.52) |
|  | MoNet | **0.61 (0.05)** | **0.57 (0.02)** |
|  | UG-SCNN | 2.03 (1.52) | 3.75 (3.17) |

**Table 9:** Predicting GA at birth, MAE in weeks on native space data. Columns refer to the presence or lack of rotational augmentations during training.The first figures in the tables represent the best MAE, with the figures in brackets representing the standard deviation.

| Method | Network | Unrotated | Rotated |
| --- | --- | --- | --- |
| Baseline | Projected ResNet | **1.49 (0.49)** | **1.49 (1.13)** |
| Spectral | S2CNN | 1.52 (0.60) | 1.67 (0.40) |
|  | ChebNet (TopK) | 1.53 (0.77) | 1.93 (0.33) |
|  | ChebNet (w/o pooling) | 1.70 (0.36) | 1.83 (0.07) |
|  | GConvNet (TopK) | 2.14 (0.34) | 2.11 (0.11) |
|  | GConvNet (w/o pooling) | 2.30 (0.74) | 1.60 (0.45) |
| Spatial | Spherical UNet | 2.16 (0.57) | 2.57 (0.02) |
|  | MoNet | 1.58 (0.06) | 1.52 (0.22) |
|  | UG-SCNN | 2.89 (2.17) | 3.29 (2.19) |

**Figure 8:**
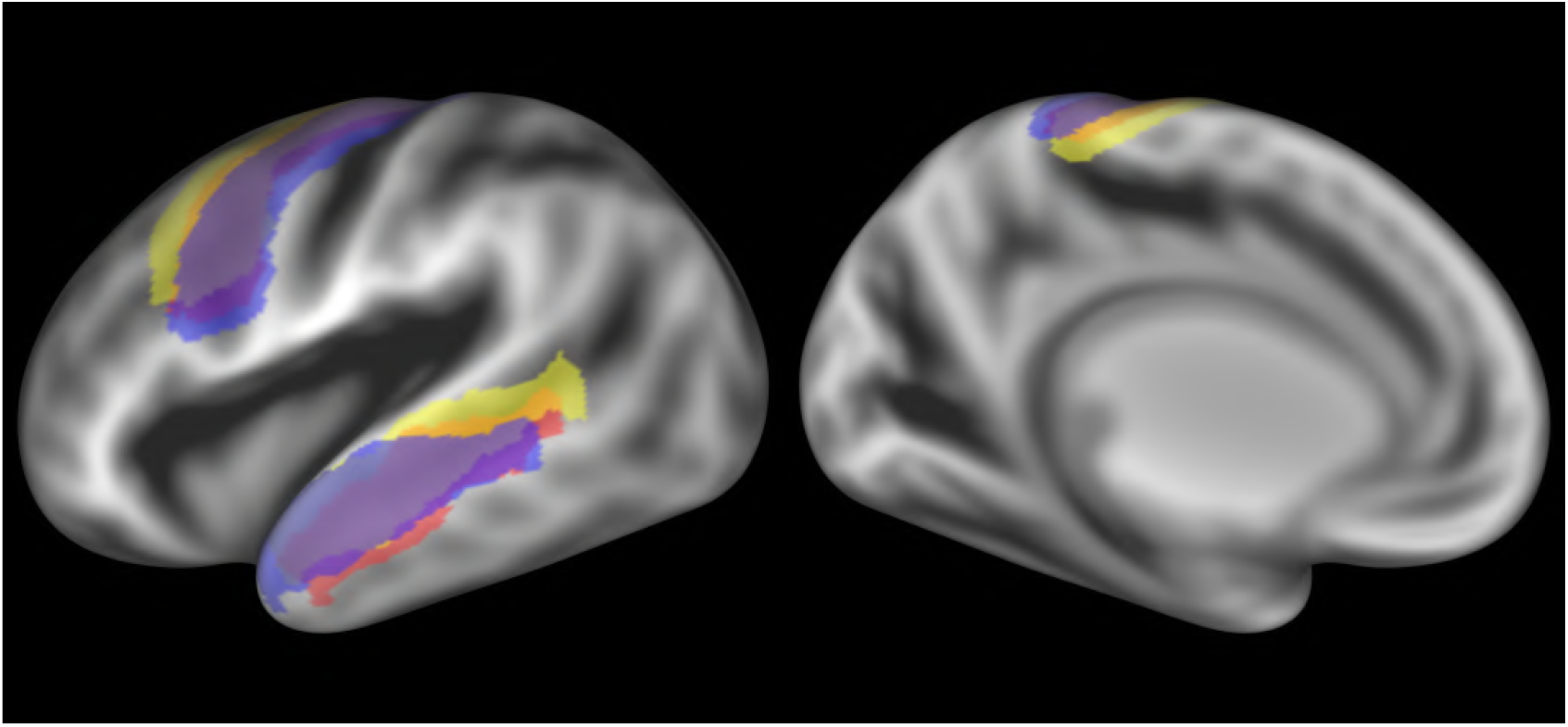
The postcentral (top) and inferior temporal (bottom) gyri labels for 3 subjects in native space (blue, red, yellow) overlaid on the 40-week sulcal depth map (grey-black) on a very inflated 40-week PMA left hemispheric surface.

The results on prediction of PMA at scan in native space are comparable to those on template space for almost all models. However, spherical UNet which was previously the best performing model in template space showed a large drop in performance in native space, rising from a MAE of 0.57 to 0.87 (when trained without rotations). This result indicates that it was less able to account for the variability of locations in native space. In general, NRE models suffered a loss in performance when rotations were included during training, whereas RE methods displayed a mix of improvements and losses. Overall, MoNet was the best performing model with a MAE of 0.610 weeks (trained without rotations) and 0.570 weeks (with rotations).

The results on the prediction of GA at birth were less consistent. Projected ResNet performed the best, with an MAE of 1.49 weeks (with and without training rotations), similar to its performance on template space. Similarly, S2CNN and MoNet, which both performed consistently well in template space, also performed well on both native experimental setups with MAEs of 1.52/1.67 and 1.58/1.52 weeks when trained without/with rotations respectively. Native space graph models continue to perform worse than more expressive models, with variable increases and decreases in performance relative to template space models. SCNN-UG remained a consistently poor performing model, returning the highest MAEs across both experiments (MAEs of 2.89/3.29 weeks). Spherical UNet again displayed consistently poor performance, returning a MAE of over 2 weeks for both experiments, consistent with its results in template space when training with rotations, and in stark contrast to its performance as the best model in template space without rotations. This highlights the inability of Spherical UNet to learn from data that is not registered to a template space. Finally, while it may appear that UG-SCNN improved for U/U on native space data, it only returned values within a narrow range around the mean; this suggests that these predictions relied heavily on the scan age confound.

The results of the segmentation experiments on native space are shown in Figure 9 and Table 10. Here, MoNet and Spherical UNet, which were the best performing methods in template space by a small margin, were also the best two performing methods in native space with Dice overlap ratios of 0.940 and 0.937 respectively, where we report results trained with rotations to account for possible memorisation witnessed in template space. These were followed by the variants of GConvNet (0.926) and ChebNet (0.924) without TopK pooling, and Projected ResNet (0.897). UG-SCNN had the steepest drop in performance from a Dice overlap ratio of 0.887 in template space to 0.769 (unrotated) in native space, which could be attributable to it’s dependence on (non-rotationally equivariant) hand-engineered features (Table 1). In general, native space segmentation performed similarly to template space, which may be partly explained by the fact that there is still significant overlap of regions - the mean Dice overlap ratio between the group average in native space and the single subject native space labels (across both hemispheres) was 0.742. Importantly, however, most methods (including Projected ResNet) outperformed this score, which suggests that the performance of these models cannot be entirely attributed to memorisation. As seen in template space, the TopK pooling-based models also failed in native space.

**Table 10:** Mean Dice overlap ratio for cortical segmentation data in native space and standard deviation across all subjects.

| Method Type | Network | Unrotated | Rotated |
| --- | --- | --- | --- |
| Baseline | Projected Residual UNet | $0.932 \pm 0.020$ | $0.897 \pm 0.071$ |
| Spectral | ChebNet (TopK) | $0.260 \pm 0.071$ | $0.255 \pm 0.077$ |
| | ChebNet (w/o TopK) | $0.924 \pm 0.014$ | $0.879 \pm 0.033$ |
| | GConvNet (TopK) | $0.238 \pm 0.074$ | $0.240 \pm 0.070$ |
| | GConvNet (w/o TopK) | $0.926 \pm 0.013$ | $0.917 \pm 0.012$ |
| Spatial | Spherical UNet | $0.944 \pm 0.010$ | $0.937 \pm 0.013$ |
|  | MoNet | **$0.945 \pm 0.008$** | **$0.940 \pm 0.010$** |
| | UG-SCNN | $0.769 \pm 0.081$ | $0.327 \pm 0.039$ |

**Figure 9:**
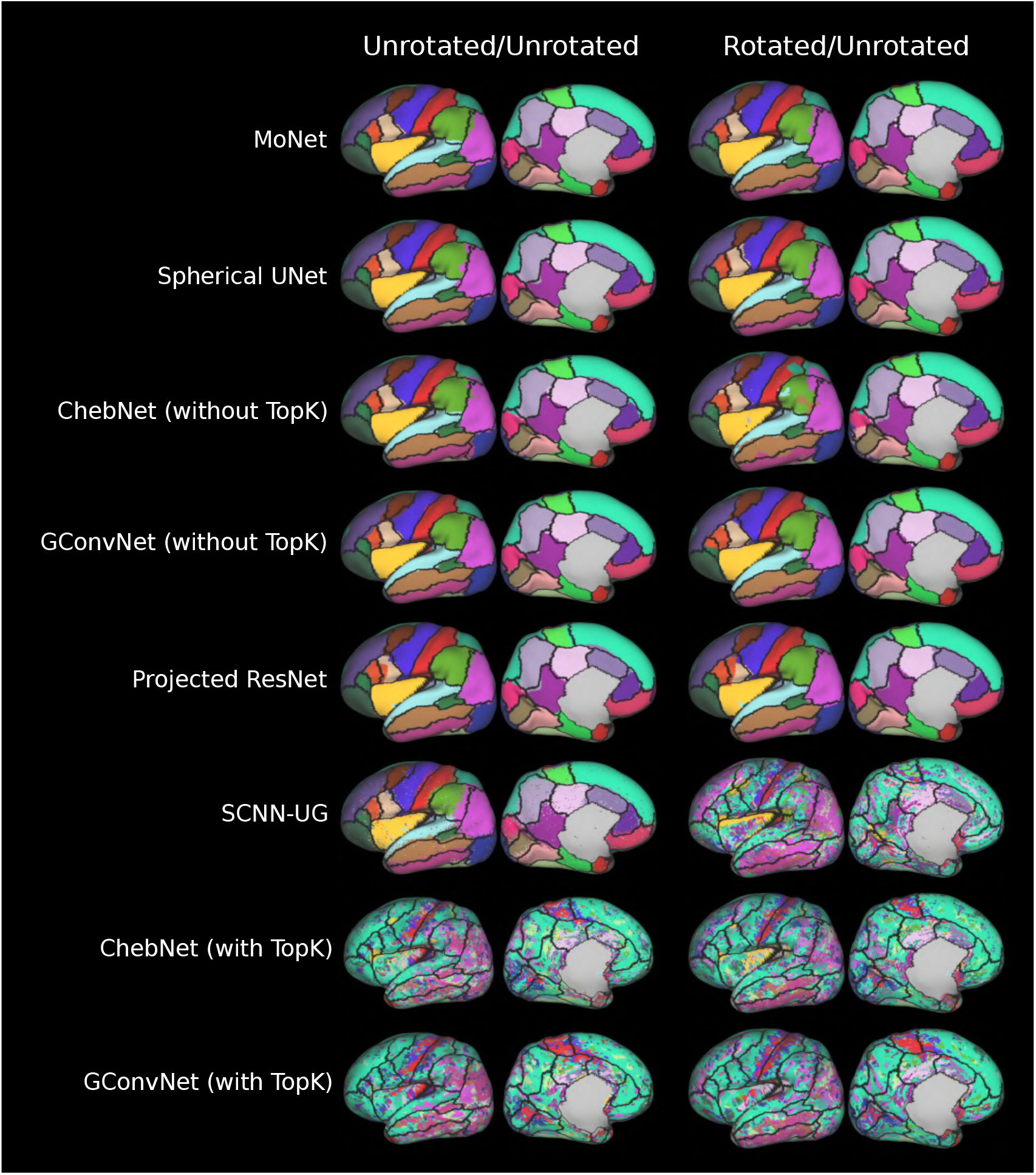
Example of predicted cortical surface segmentations for a single subject across all models for U/U and R/U experiments in native space. Black lines show ground truth labels from M-CRIB-S pipeline. Results are shown on a subject-specific very inflated left hemispheric surface.

### 4.5. Comparison of Learning on Different Structures

Results for comparisons against PointNet++ and Spectral Embedding GCN are presented in Table 11. Results show that both methods marginally outperformed the best spherical model, Spherical UNet, in predicting PMA at scan. However, they under-performed against Spherical UNet when predicting GA at birth, by a greater margin. These results are not entirely surprising, as the cortex undergoes marked morphological and microstructural changes during the perinatal period (Alexander et al., 2019a; Thompson et al., 2019). On the other hand, preterm birth affects both cortical morphology and microstructural development (Ajayi-Obe et al., 2000; Engelhardt et al., 2015; Dimitrova et al., 2021a, 2020), which makes prediction of GA at birth a harder task. These findings suggest that cortical shape alone is not sufficient for accurate phenotype prediction (Vosylius et al., 2020), and that cortical shape is not strictly necessary to accurately predict these phenotypes.

**Table 11:**
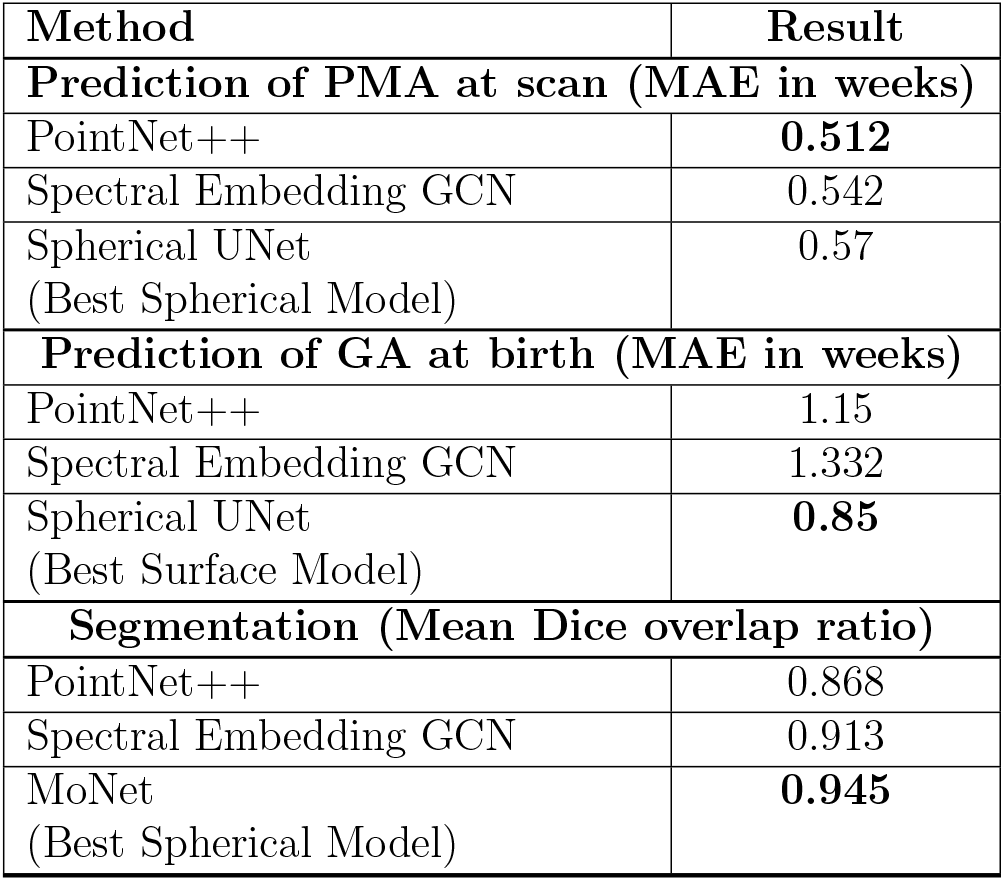
A comparison of surface deep learning models on the same tasks, but using different forms of the same cortical surface data. PointNet++ utilises the native mesh as a point cloud, Spectral Embedding GCN uses a graph spectral embedding of the native meshes and the Spherical Models use shape-agnostic icospheres.

On segmentation, both PointNet++ and the Spectral Embedding GCN performed well (Dice Overlap Ratios of 0.868 and 0.913 respectively), but under-performed relative to the spherical methods (0.945). For Spectral Embedding GCN this may be explained by its choice to embed all data into a very low (3) dimensional spectral embedding space, which may not capture all the high-frequency folding information needed for tuning group average segmentations for individual brains. Direct comparison of the spectral and Euclidean coordinate representations confirms that clear differences exist. We note that there are variations in model architecture, specifically the size of the kernels, that may also explain this models’ performance. PointNet-based networks use shared multi-layer perceptrons (MLP) or 1×1 convolutions, which retain only the most significant activation on features. This can lead to missing some detailed information for the segmentation task (Zhang et al., 2020).

### 4.6. Visualisation

Results of occlusion-based visualisation, for predicting PMA at scan, are shown in Figures 10 and 11, split by lateral and medial views, respectively. The subject used for all models was a term neonate born at 41.14 weeks GA and scanned at 43.57 weeks PMA. The images in the first column are from the simultaneous occlusion of all the modalities, and the proceeding four columns show occlusions based on modality-specific occlusion of T1w/T2w ratio myelin map, curvature, cortical thickness and sulcal depth, respectively. Figure 12 visualises occlusion in volumetric space for the 3D CNN model on the same subject and task - demonstrating at a glance that identifying cortical features is significantly easier on surface-based models. Note particularly that some regions identified on the volumetric images are not even within the subject’s brain at all - an artefact of the 3D convolutions’ receptive fields overlapping with the edge of the brain. Artefacts are also present on the surface, due to the sensitivity of all models to focus on distortions arising from the inflation process that turns the white matter surface into an icosphere. These vary with brain size (and thus gestational age) and are most prominent around the hemispheric cut on the medial aspect of the cortical surface.

**Figure 10:**
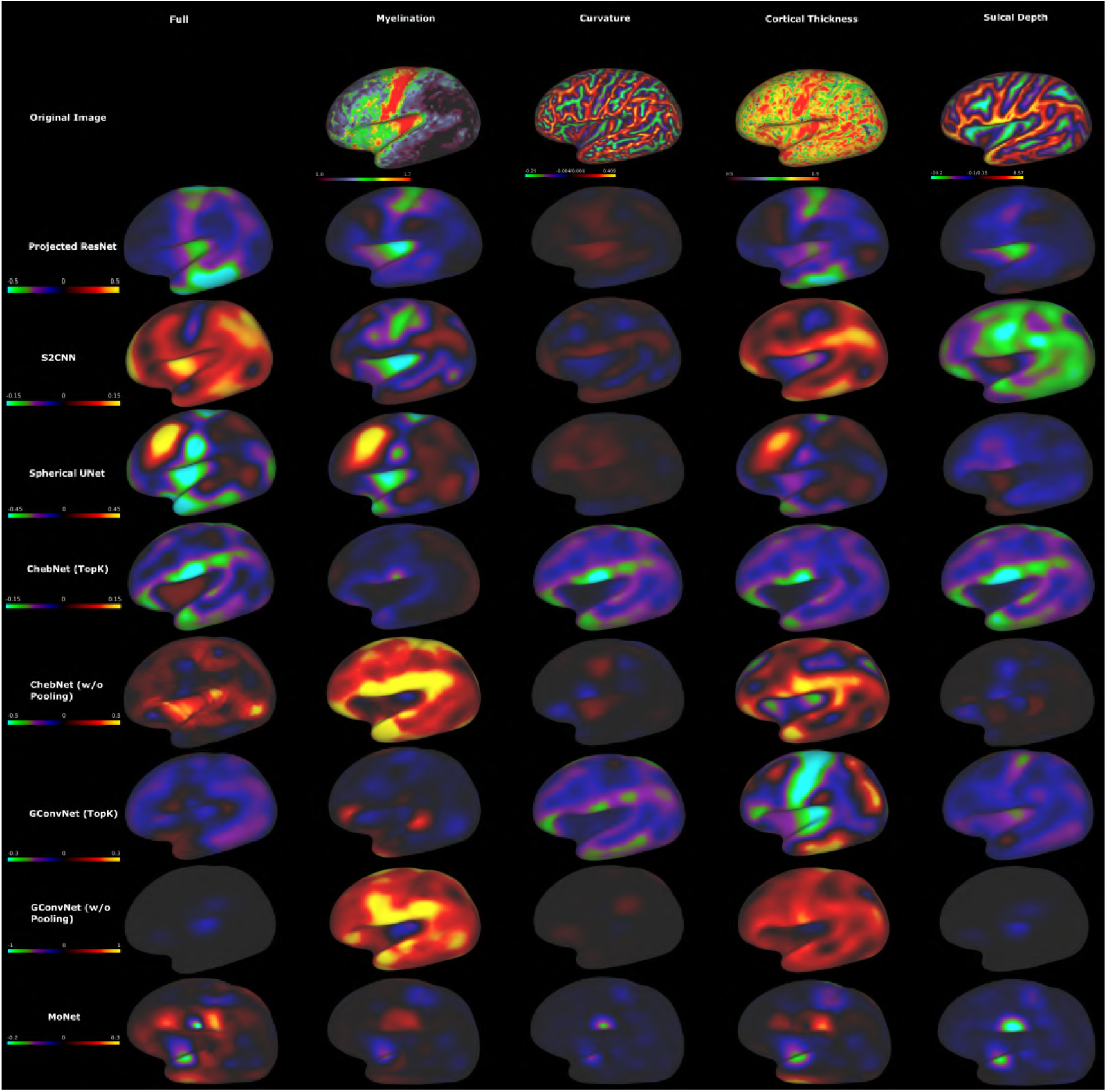
Lateral view of every surface models’ visualisation on the estimation of PMA of a term neonate born at 41 weeks and scanned at 43 weeks.

**Figure 11:**
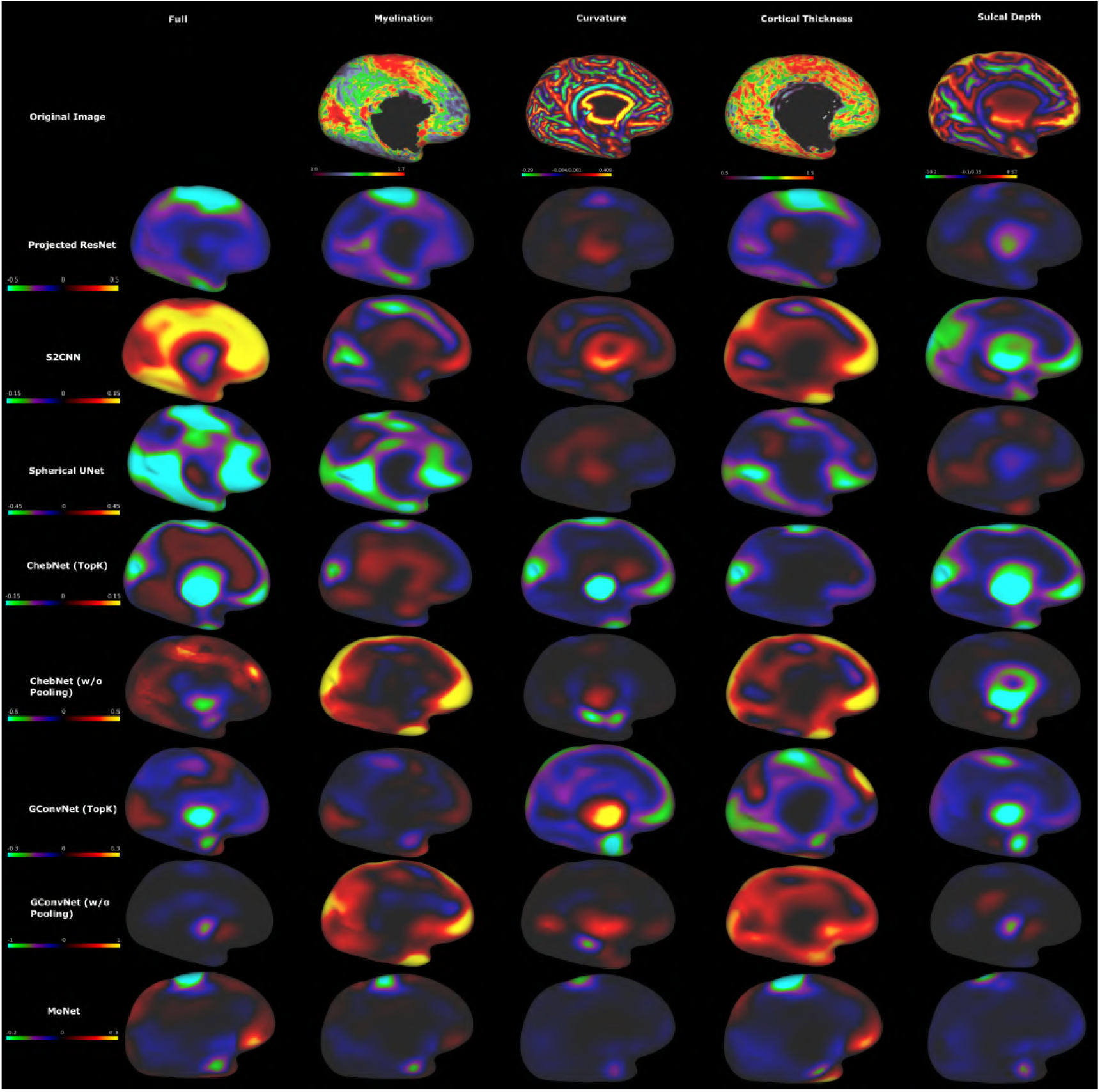
Medial view of every surface models’ visualisation on the estimation of PMA of a term neonate born at 41 weeks and scanned at 43 weeks.

**Figure 12:**
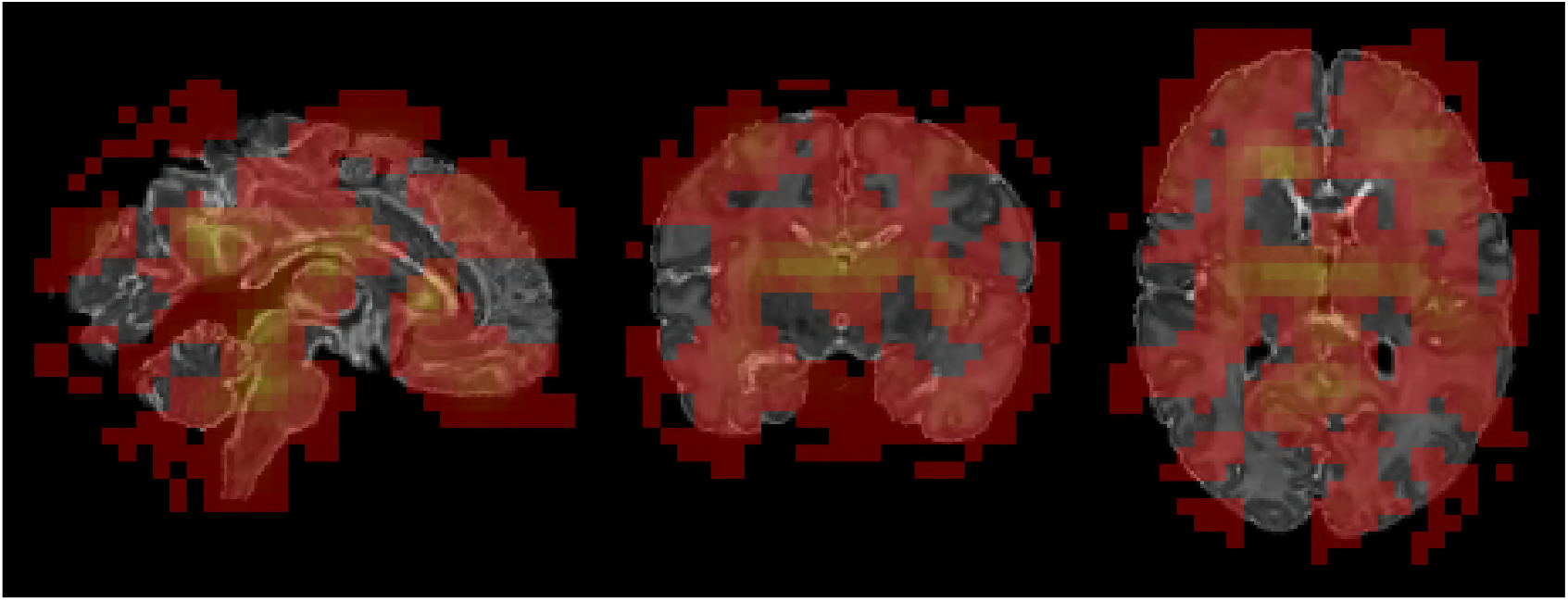
Occlusion based visualisation of the 3D Residual CNN for the estimation of PMA of a term neonate with true PMA of 43 weeks.

It can be seen from the relative intensities of the visualisations that some modalities had a greater impact on a model’s prediction than others, although not all models relied on the same modalities. It can also be seen that occluding all modalities simultaneously returned some higher errors in the resulting predictions, but not necessarily more informative visualisations. The overall sensitivity of the output predictions was variable for different models, with S2CNN and ChebNet varying the least on average, and GConvNet varying by up to a week.

Many of the visualisations were consistent across models, and correlated both with features in the original image, and known neurodevelopmental changes in the perinatal period, reinforcing the validity of the features highlighted. A notable example is the continued myelination of the central and lateral sulci on the lateral hemispheric surface, and the calcarine and parieto-occipital sulci on the medial hemispheric surface - regions that are highlighted most prominently by occlusion of the T1w/T2w ratio in S2CNN, Projected ResNet and Spherical UNet. S2CNN in particular highlighted a region in the approximate location of the middle temporal (MT)/medial superior temporal (MST) areas (Glasser et al., 2016), despite its small size. The development of cortical thickness also follows a specific spatiotemporal sequence, which was almost the entire focus of the GConvNet model.

## 5. Conclusion and Further Work

The aim of this paper was to explore the performance and future potential of geometric deep learning for registration-independent cortical surface analysis. Surface deep learning is a broad field and encapsulates a variety of extensions to the traditional Euclidean convolution. Extending the convolution operation to surfaces is non-trivial and existing methods incorporate some form of compromise: either in computational expense, expressive power or rotational equivariance. The results of this study indicate that the optimal balance of these properties are task and data-dependent.

Reassuringly, the observed degrees of rotational equivariance matched very well with prediction, where RE models generalised well to unseen rotations but the NRE models did not. Moreover, this contrast could not be bridged by simply including rotational augmentations during training. And, this same pattern was observed for experiments trained on unregistered data. Overall the optimal balance between rotational equivariance and filter expressivity seemed to be most specific to task and, to a lesser extent, space (template vs. native). For the prediction of age phenotypes in template space, filter expressivity was much more important than rotational equivariance - with S2CNN and Spherical UNet, the methods that prioritise filter expressivity, outperforming all the other methods that sacrificed these for computational efficiency, provided the data was pre-aligned. On segmentation, the results were more mixed, with no clear advantage of either rotational equivariance nor filter expressivity. Instead, it seems that the method of pooling holds greater importance, as the models that used TopK pooling successfully on the prediction of age phenotypes, completely failed on segmentation tasks.

Importantly, comparing the performance of surface deep learning techniques relative to non-surface baselines, it was observed that, on phenotype prediction, the 3D CNN trained on the volumetric data did not outperform most of the models that were trained exclusively on cortical surface data. This is a strong indication that surface-based features are sufficient to learn these phenotypes. Further, the visualisations produced in the volume were significantly less informative for identifying cortical features than those produced directly on the surface. Surface deep learning techniques also reliably outperformed simple ROI analysis on all tasks. They generally outperformed Projected ResNet on cortical segmentation, and with respect to transformation equivariance; although, Projected ResNet performed well on phenotype prediction tasks.

Experiments benchmarking spherical methods against methods that prioritised learning descriptors of cortical shape properties (PointNet++ and Spectral-Embedding GCN) demonstrated that, while shape was clearly useful, particularly for scan age prediction, it was not necessary for this task. Poorer performance of these methods (relative to spherical networks) on birth age may reflect the fact that preterm birth is known to affect cortical morphology (Engelhardt et al., 2015; Makropoulos et al., 2016), even in the absence of differences in cortical surface area or volume (Dubois et al., 2019). However, changes in cortical shape are known to be strong features of cortical growth, in both health (Alexander et al., 2019a; Thompson et al., 2019) and disease.

These results provide a clear strategy for the future development of an optimal model for deep learning on the cortical surface. They highlight that the type of task is the most crucial factor in prioritising the properties of a surface-based model, and suggest a difference between learning on a global context, such as predicting a phenotype, and a local context like segmenting a cortical region. The former requires expressive filters capable of learning complex features, the latter requires models capable of capturing the full spatial context on a sphere. Both suggest that an optimal model would have all three of the desired properties: rotational equivariance, low computational overhead and high expressive power. This fits our findings that the best model overall of those benchmarked was MoNet, which was found to be relatively inexpensive, expressive and rotationally equivariant. One approach for a more powerful future model would be extending existing architectures appropriately. For example, adapting 2D ResNets, trained on flattened cortical data, to include rotational equivariance, improving the filter expressivity of graph-based methods, or reducing the large computational overhead of S2CNN, which performed consistently across all tasks in both native and template space but was constrained by its large computational cost. Aligned with this observation, a recent extension to Spherical UNet has been published which trains deformable hexagonal filters to allow for greater flexibility during segmentation (Zhao et al., 2021).

An interesting extension to this work would be to apply the models to the significantly more complex task of multimodal parcellation of adult cortical surfaces from the Human Connectome Project (HCP) (Glasser et al., 2016). This task infers functional organisation of the cortex, with very limited correlation with cortical shape. Preliminary work on using an ensemble of gDL networks to solve this problem was presented in Williams et al. (2021a). At the same time, greater benefit for registration independent surface deep learning techniques may be seen through applying the models to learning more complex and heterogeneous phenotoypes, such as predicting neurodevelopmental outcomes, or cognitive test scores.

While occlusion-based visualisation provides some evidence that surface deep-learning techniques could provide interpretable, registration independent frameworks for investigating cortical organisation, saliency-based methods for interpreting deep networks are limited in the sense that they focus only on the most prominent and consistent features required for the prediction of task (Bass et al., 2020, 2021; Baumgartner et al., 2018). Future work may benefit from taking inspiration from the field of interpretable and explainable machine learning, to generate more holistic models of cognition and disease, for example: using techniques for image-to-image translation (Bass et al., 2020, 2021; Baumgartner et al., 2018; Schutte et al., 2021) to discover all features required to translate an image from one class to another, or incorporating ideas and concepts from the causal modelling domain (Pawlowski et al., 2020). The data and benchmarks established and made available in this work provide a framework for further investigation in this topic.

## 6. Acknowledgements

The authors would like to thank the participants and families recruited to the dHCP, and all the neonatal staff at the Evelina Newborn Imaging Center, St. Thomas’ Hospital, Guy’s & St. Thomas’ NHS Foundation Trust, London, UK. The dHCP project was funded by the European Research Council (ERC) under the European Union Seventh Framework Programme [FR/2007–2013]/ERC Grant Agreement no. 319,456. This study was supported in part by the Wellcome Engineering and Physical Sciences Research Council Centre for Medical Engineering at King’s College London [grant WT 203,148/Z/16/Z] and the Medical Research Council (UK) [grant MR/K006355/1]. A.F is supported by an Engineering and Physical Sciences Research Council (EPSRC) United Kingdom Doctoral Training Partnership (DTP) - EPSRC DTP 2018 [EP/R513064/1]. L.Z.J.W is supported by funding from the Commonwealth Scholarship Commission, United Kingdom. E.C.R is supported by an Academy of Medical Sciences/the British Heart Foundation/the Government Department of Business, Energy and Industrial Strategy/the Wellcome Trust Springboard Award [SBF003/1116] and Wellcome Collaborative Award [215573/Z/19/Z]. A.D.E. receives support from the Medical Research Council Centre for Neurodevelopmental Disorders, King’s College London [grant MR/N026063/1]. G.B. was supported by a National Health and Medical Research Council (Australia) Investigator Grant [1194497].

The authors acknowledge use of the research computing facility at King’s College London, Rosalind (https://rosalind.kcl.ac.uk), which is delivered in partnership with the National Institute for Health Research (NIHR) Biomedical Research Centres at South London & Maudsley and Guy’s & St. Thomas’ NHS Foundation Trusts, and part-funded by capital equipment grants from the Maudsley Charity (award 980) and Guy’s & St. Thomas’ Charity (TR130505). The views expressed are those of the author(s) and not necessarily those of the NHS, the NIHR, King’s College London, or the Department of Health and Social Care.

## Appendix A. Convolution in the Spectral and Spatial Domains

A convolution is an operation that combines two signals to produce a third signal. In deep learning, one of these is a static input signal from the data and the second is a variable filter. The training process ‘learns’ useful filters that can extract information from the signal to complete the task. The convolution theorem states that for two signals, *f* and *g*, the following is true - for any geometry - including spheres:

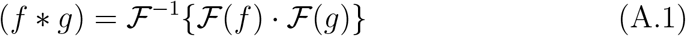

Here * denotes a convolution, · is a simple multiplication and *ℱ, ℱ*^−1^ are Fourier and inverse Fourier transforms respectively.

In a general sense, a Fourier transformation decomposes a signal into a weighted sum of orthogonal components (or spectra). Equation A.1 should therefore be seen as defining the convolution in one of two different ways: the *spatial* approach (represented by the left hand side of the equation), which involves directly evaluating a discrete integral over the two functions, and the *spectral approach* (represented by the right hand side of the equation), which offers an alternative formulation that relies only on defining some (domain-dependent) Fourier transform over the chosen geometrical space.

### Appendix A.0.1. Spatial Methods

The convention for Euclidean domains is to implement spatial convolutions as a translation of an filter (*g*) across an image (*f*):

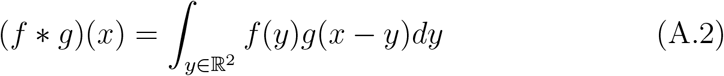

Where this is calculated as the integral of the product of the image and filter, at each position *x*, as the filter is translated across the image. In the finite discretised (pixelated) space of actual Euclidean data, the integral reduces to a discrete sum, and the shifts *y* are restricted only to the subspace of the Euclidean domain where the filter and the image is overlap. This explains the common interpretation of Euclidean convolutions as the filter ‘sliding’ along the image.

Spherical convolutions are similarly defined as the overlap between a filter over an image, but the relative transformations of the two are not translations but rotations along the surface of the sphere, given by a rotation matrix *R*. The integral operation, when defined on the sphere, becomes a surface integral (Cohen et al., 2018):

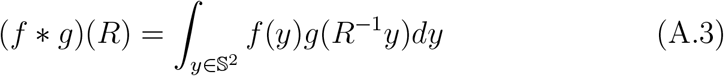

where *R* is an element of the spherical rotation group (Rotman, 1995) and the operation is integrated over all possible rotations of the (spherical) filter *g* relative to the spherical signal *f*. Here, *dy* is redefined as the standard integration measure on the sphere.

Due to the complexity of evaluating this integral directly, most spatial methods instead work by implementing a ‘template-matching’ approach, which for Euclidean convolutions is implemented not by sliding a filter over the image, but rather by splitting up the image into patches (the same shape and size as the filters). In this way, the 2D/3D convolutional operation can be efficiently, and correctly, implemented as matrix multiplication - a linear transform (**Z** = **WF**) of kernel weights with the image features; where, the rows of **W** represent filters and the columns of **F** represent patches of the image (or activation maps).

For surfaces, the most direct translation of this operation is implemented by fitting filters of a certain regular shape and tessellation (e.g. hexagon or square, Figure A.1a) to each vertex on the manifold. The problem with this approach is that for a sphere, or general manifold, there is no fixed definition of a consistent coordinate system; thus, sliding a filter along different paths on the sphere would result in an inconsistent definition of filter orientation (Figure A.1b). As such, this means filters estimated in this way, are statically estimated and do not approximate any transformation over the surface, meaning that, while computation is fast and the features learnt may be expressive, the network cannot be rotationally equivariant; thus features will be location (and thus registration) dependent.

### Appendix A.0.2. Spectral Methods

Spectral convolutions represent the right hand side of the convolution theorem (eq. (A.1)), and their implementation varies according to what choice of domain-dependent generalised Fourier transform is used to decompose the data. For spheres, this is represented by the well known spherical harmonics basis. However, one important detail of this operation is that the output domain is no longer the sphere but rather a general 3D manifold SO(3) (representing the space of all 3D rotations).

**Figure A.1:**
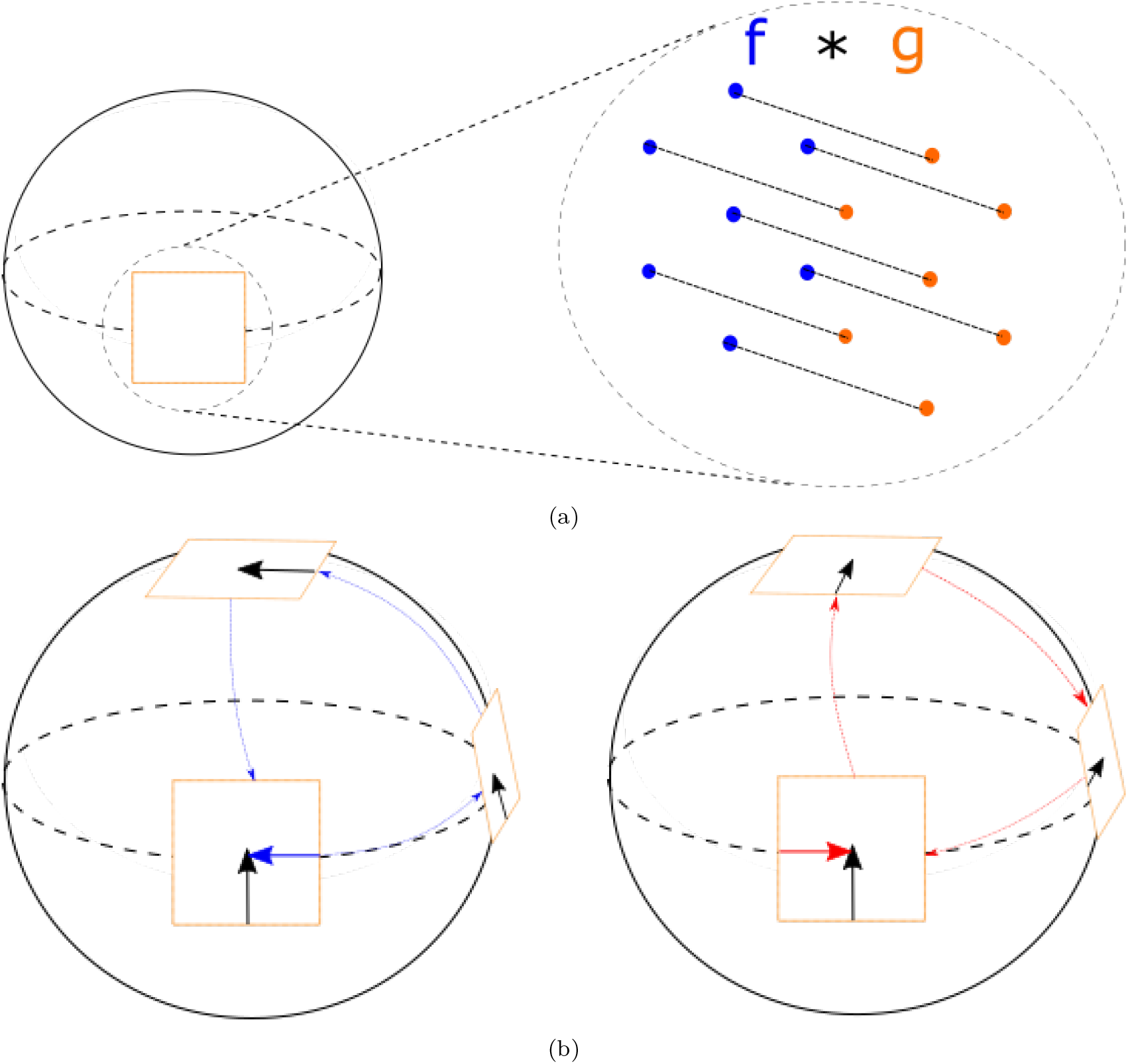
(a) Template Matching of a filter on the surface of a sphere is to multiply elementwise the (in this case hexagonal) filter (orange) values with the corresponding surface points (black) on the sphere. (b) On the sphere, filter orientation is path dependent as it moves around the surface. This can be seen from the differing ending orientations of a filter (orange) rotated along the sphere in one path (blue) compared to a different path (red).

Two approaches have been proposed to resolve this: Esteves et al. (2018), which implements all convolutions using spherical harmonic transforms, thereby keeping the output of the convolution on the sphere, but restricting the expressivity of learnable spherical filters to be rotationally symmetric about the z axis; and S2CNN (Cohen et al., 2018) (benchmarked here), which follows initial *S*^2^ convolution with a similar spectral convolution in SO(3) based on Wigner D-matrices (Martin, 1960). Such filters are more expressive, and fully rotationally equivariant; however, the extra dimension makes S2CNN very computationally expensive.

Fourier transforms may also be defined on general graphs, and this is the basis of spectral graph-based methods. The graph Fourier transform is based on an eigendecomposition of the graph Laplacian: **L** ≡ **D-A** (for an unweighted graph). Here **D, A** are the degree matrix and adjacency matrix respectively. **A** is an *N* × *N* binary adjacency matrix, representative interchangeably of an abstract graph, or mesh with vertices *v*_*i*_:

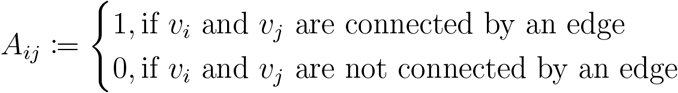

and **D** (*d*_*ii*_ = Σ_*j*_ *A*_*ij*_) is the (diagonal) degree matrix. Defined in this way, the convolution has no sense of spatial locality and a very large number of parameters. Constructing filters by adding weighted polynomial powers of the Laplacian resolves both these issues, with polynomial orders up to *k* allowing filter sizes of up to *k* hops on the graph. Here, 1 hop represents each node’s direct neighbours and 2 hops represents the next nearest neighbours (and so on). However, as it is desirable to avoid expensive repeated computation of the eigenvectors of **L**, methods typically circumvent the full calculation and replace it with a polynomial approximation (Defferrard et al., 2016; Kipf and Welling, 2017).

### Appendix A.0.3. Graph Pooling

Graph convolutional networks, by their nature, do not put any restrictions on the shape of the images, whether spherical, square or otherwise. This allows for intelligent graph coarsening techniques such as TopK pooling (Gao and Ji, 2019), which iteratively remove graph nodes based on removing the least informative nodes. We choose to benchmark the performances of ChebNet and GConvNet both with and without TopK pooling. While, the rotational equivariance of these graph convolutions depends on the chosen sampling of the sphere (Perraudin et al., 2019), for our chosen sampling scheme, meaningful rotational equivariance by both ChebNet and GConvNet is expected (Yang et al., 2020).

## Footnotes

1 http://www.developingconnectome.org

2 https://github.com/BioMedIA/dhcp-structural-pipeline

3 https://www.github.com/DevelopmentalImagingMCRI/MCRIBS

4 https://github.com/ecr05/Random_forests_for_cortical_imaging_data

5 Adapted from https://github.com/amiralansary/BrainSurfaceTK

6 https://github.com/ecr05/Random_forests_for_cortical_imaging_data

7 https://github.com/jonas-koehler/s2cnn

8 https://github.com/maxjiang93/ugscnn

9 https://github.com/zhaofenqiang/Spherical_U-Net

10 https://github.com/kharitz/spectral_embedding_GCN

11 from https://biomedia.github.io/dHCP-release-notes/download.html

12 https://gin.g-node.org/lzjwilliams/geometric-deep-learning-benchmarking/

13 https://github.com/Abdulah-Fawaz/Benchmarking-Surface-DL

